# USP19 modulates cancer cell migration and invasion and acts as a novel prognostic marker in patients with early breast cancer

**DOI:** 10.1101/2020.07.01.181883

**Authors:** Fabiana A Rossi, Juliana H Enriqué Steinberg, Ezequiel H Calvo Roitberg, Molishree Joshi, Ahwan Pandey, Martin C Abba, Beatrice Dufrusine, Simonetta Buglioni, Vincenzo De Laurenzi, Gianluca Sala, Rossano Lattanzio, Joaquín M Espinosa, Mario Rossi

## Abstract

Tumor cell dissemination in cancer patients is associated with a significant reduction in their survival and quality of life. The ubiquitination pathway plays a fundamental role in the maintenance of protein homeostasis both in normal and stressed conditions and its dysregulation has been associated with malignant transformation and invasive potential of tumor cells, thus highlighting its value as a potential therapeutic target. In order to identify novel molecular targets of tumor cell migration and invasion we performed a genetic screen with an shRNA library against ubiquitination pathway-related genes. To this end, we set up a protocol to specifically enrich positive migration regulator candidates. We identified the deubiquitinase USP19 and demonstrated that its silencing reduces the migratory and invasive potential of highly invasive breast cancer cell lines. We extended our investigation *in vivo* and confirmed that mice injected with USP19 depleted cells display increased tumor-free survival, as well as a delay in the onset of the tumor formation and a significant reduction in the appearance of metastatic foci, indicating that tumor cell invasion and dissemination is impaired. In contrast, overexpression of USP19 increased cell invasiveness both *in vitro* and *in vivo*, further validating our findings. More importantly, we demonstrated that USP19 catalytic activity is important for the control of tumor cell migration and invasion, and that its molecular mechanism of action involves LRP6, a Wnt co-receptor. Finally, we showed that USP19 overexpression is a surrogate prognostic marker of distant relapse in patients with early breast cancer. Altogether, these findings demonstrate that USP19 might represent a novel therapeutic target in breast cancer.

## INTRODUCTION

Cell migration plays a crucial role in a wide variety of physiological processes such as development, tissue injury and wound healing [1]. Its activation is highly regulated both spatially and temporarily, contributing to the maintenance of tissue and cellular homeostasis [1, 2]. Therefore, it is not surprising that when deregulated, migration is associated with the development and progress of multiple pathologies, including cancer [2–4].

During the development of malignant tumors, transformed cells change and acquire the ability to invade and abandon their original position. In order to do so, they need to pierce the surrounding extracellular matrix (invasion) and reach the circulatory torrent (intravasation). If they survive, some will be able to disseminate and get through distant capillary walls (extravasation), to invade the extracellular matrix in a new host environment, establishing a secondary tumor (colonization). During this process, cells acquire the ability to proliferate in an anchorage-independent manner with elevated invasive potential [5]. Cell motility plays a vital role in many of these events and therefore has a major influence in metastatic cell dissemination. Particularly in cancer, alteration or exacerbation of malignant tumor cell migration and dissemination is the principal cause of death due to solid tumors [6].

In addition, it was observed that decreasing the migratory capabilities of tumor cells can restore a certain level of sensitivity to cytotoxic reagents and increase the susceptibility to chemotherapeutic treatments [7, 8]. Consequently, targeting genes that regulate cell motility could be beneficial in the treatment of highly aggressive cancers [9, 10], and anti-migratory or anti-invasive activity is usually viewed as a desired attribute for novel anticancer drugs [11].

Cell motility is a complex process that requires post-translational regulation of wide variety of proteins, modifying their biological function, subcellular localization, or half-life. Ubiquitination is an important form of protein post-translational modification that consists in the conjugation of ubiquitin polypeptides to target proteins [12, 13]. Ubiquitin tagged proteins are either subjected to destruction or responsible for regulating different processes, including endocytosis, DNA repair, cell cycle regulation, and gene expression [14]. The process of ubiquitin chain conjugation and elongation are regulated by a complex enzymatic cascade [15]. Also, like most post-translational modifications, the addition of poly-ubiquitin chains can be reversed or modified. This process is carried out by deubiquitinating enzymes (DUBs), a family of proteins including approximately 100 members classified into few sub-groups according to their catalytic core domains [16]. The ubiquitination cascade is of vital importance for the maintenance of cellular homeostasis by regulating a wide variety of processes, including cell migration and invasion [14, 17].

Therefore, in order to identify novel molecular targets within the ubiquitination pathway that positively regulate migration we conducted a loss-of-function genetic screen. Screens targeting multiple genes significantly shortens experiment duration [18], and therefore they have been widely and successfully used to find genes involved in many physiological and pathological cellular processes [18–22], as they are relatively easy to perform and implement. In this study, we used a pooled shRNA interference library and an immortalized tumorigenic mammary epithelial cell line, MDAMB231, derived from a human triple-negative breast cancer. This type of cancer lacks expression of the estrogen receptor, progesterone receptor and human epidermal growth factor receptor 2 (HER2) and is associated with aggressive behavior and an overall poor prognosis [23]. Currently, there are no specific targeted therapeutic systemic alternatives for its treatment [24–26], and optimal chemotherapy regimens have yet to be established.

From our screen, we identified the Ubiquitin specific protease 19 (USP19) as a new candidate gene associated with the regulation of cell migration and invasion. USP19 presents different isoforms and the most distinctive feature, structurally and functionally, is that some of them have a cytoplasmic localization while others have a transmembrane domain that serves as anchorage to the endoplasmic reticulum [27, 28]. This DUB is associated with protein quality control and cellular homeostasis [27–31]. In particular, it has been demonstrated that USP19 regulates LRP6 stability, a co-receptor of the Wnt signaling cascade [32]. Aberrant activation of this pathway and LRP6 polymorphisms and overexpression have been associated with susceptibility to the development of different cancers, including breast cancer [33–37].

To validate USP19 function as a positive regulator of migration and invasion, we performed a series of *in vitro* and *in vivo* experiments analyzing USP19’s role in colonization and tumor formation. In addition, we showed that USP19 overexpression is associated with distant relapse in patients diagnosed with early breast cancer (T1-2, N0, M0). Collectively, our data suggest that USP19 plays a crucial role in breast cancer cell dissemination, and we provide novel evidence that it can be a prognostic marker and attractive candidate for the development of new therapeutic strategies in breast cancer.

## RESULTS

### Migration-based screen to identify ubiquitination pathway genes with novel regulatory functions

In order to identify novel positive regulators of cell migration within the ubiquitination pathway, we performed an shRNA-based functional selection screen (Fig. 1A). A pooled recombinant lentiviral shRNA library targeting over 400 human ubiquitination related genes (≈ 5 shRNAs per gene) was stably transduced into breast cancer cells. The functional selection consisted in placing the mixed population into the upper compartment of a transwell unit and allowing migration through the perforated membrane to the lower compartment. Cells that exhibited reduced migration were isolated from the upper compartment and amplified. We performed subsequent enrichment cycles until shRNA-transduced cells lost about 80% of their initial migratory potential (Fig. 1B). After every enrichment cycle, we evaluated shRNAs relative abundance in the cell population by PCR amplification and quantitative sequencing from genomic DNA. As shown in Figure 1C, as enrichment cycles increased, we observed a marked reduction in the number of shRNAs, suggesting that the selection process was efficient. As a control, we used an empty vector-transduced cell line.

**Figure 1.**
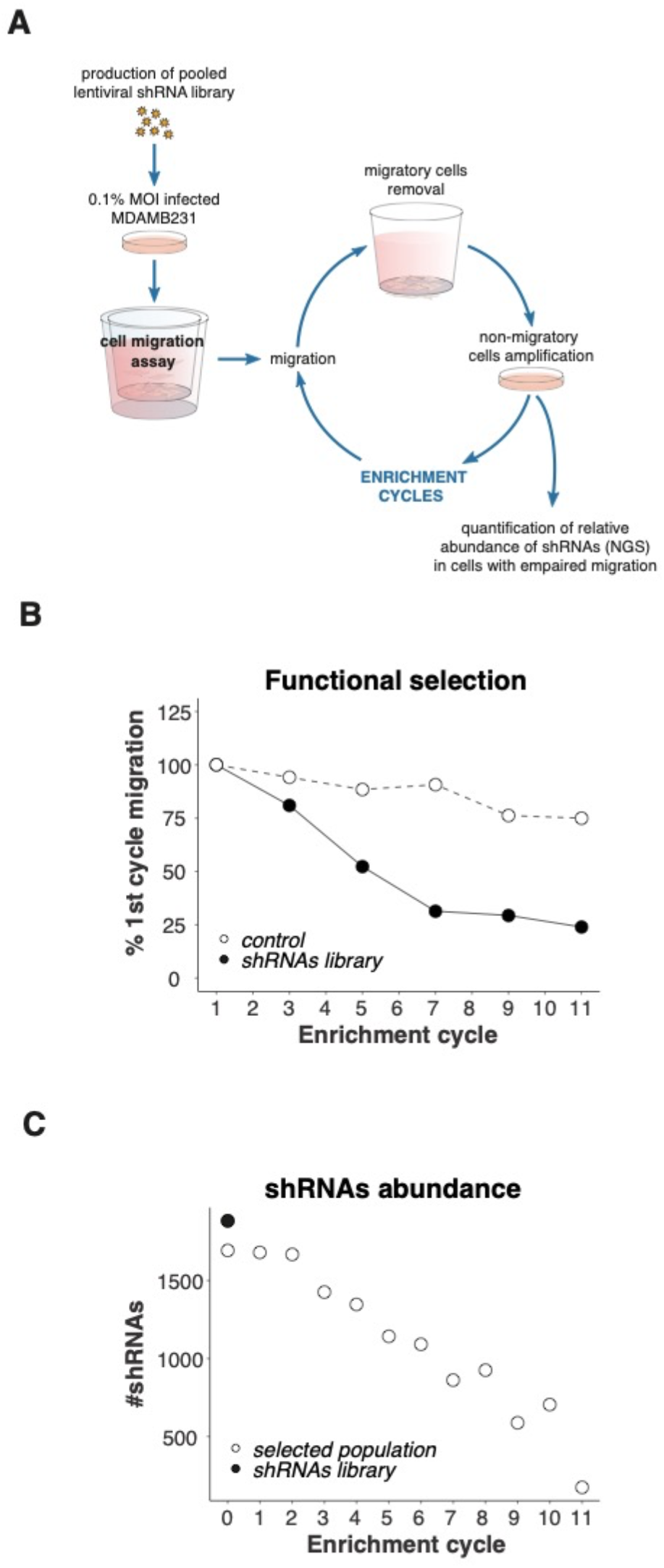
shRNA-based selection of positive regulators of cell migration. (**A**) Overview of the selection procedure. The production and infection of a ubiquitination-related lentiviral shRNA library are described in Methods. Two weeks after lentiviral infection and selection, MDAMB231 cells were seeded onto transwell inserts and allowed to migrate across the porous membrane for 24 hours in order to select cells with a decreased migration phenotype. Migrating cells were removed and non-migrating cells were collected from the inserts upper compartment and amplified. Cells were then reseeded onto transwell culture inserts for a subsequent cycle of selection; this procedure was repeated until cells lost 80% of their initial migratory potential. After every cycle of selection, the relative abundance of the different shRNAs was evaluated using Next Generation Sequencing. (**B**) Transwell assay was used every other enrichment cycle to determine the percentage of migratory cells and monitor the selection process. (**C**) shRNAs’ abundance was estimated after each selection cycle.

### Selection of candidate genes

After the selection process, we followed an analytical workflow to select candidate genes for further validation (Fig. 2A). In order to avoid false positives due to off-target effects, we discarded those genes for which only one shRNA targeting its sequence was found in the sequencing results. These criteria allowed us to identify 30 genes whose depletion altered migration. Half of these genes had already been associated with migration, invasion, metastasis or tumorigenesis, and served as a proof of principle for the efficacy and specificity of our screen (Supp. Fig. 1 and Supp. Table 1). Among the identified candidates, we focused our attention on the study of the deubiquitinase USP19.

**Figure 2.**
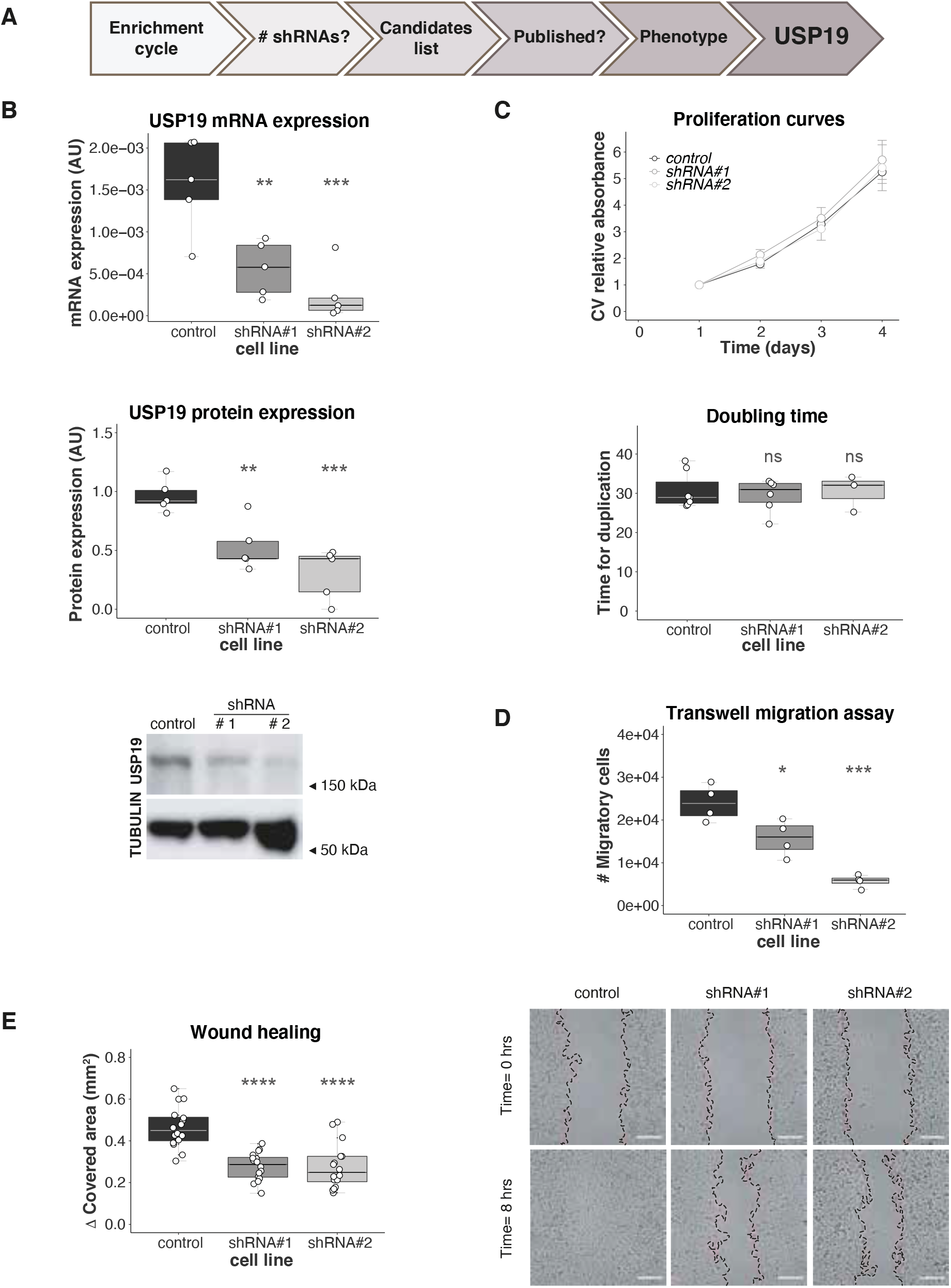
Validation and characterization of USP19 as a regulator of cell migration. (**A**) Workflow used to select a candidate regulatory gene. MDAMB231 cells were stably transduced with control empty vector shRNA (control) or two different shRNAs (#1 & #2) targeting USP19. (**B**) Efficiencies of shRNA-mediated knockdown were confirmed by RT-PCR (top, n= 5, one-way ANOVA, Dunnett’s multiple comparison test. shRNA#1 p= 0.0048 and shRNA#2 p= 0.0006) and Western Blotting (middle and bottom, n= 5, one-way ANOVA, Dunnett’s multiple comparison test. shRNA#1 p= 0.0064 and shRNA#2 p= 0.0003). (**C**) Crystal violet (CV) staining was used to determine cell growth over time. Cells were seeded onto wells and allowed to attach. At the indicated time points, cells were fixed and then stained at the end of the experiment. The graph on the top shows the mean relative CV absorbance every 24 hours. Doubling time was calculated for control and USP19 silenced cell lines on the bottom (n≥ 3, Kruskal-Wallis, Dunn’s multiple comparison test. shRNA#1 p> 0.9999 and shRNA#2 p> 0.9999). Migratory potential was evaluated by two different experiments. (**D**) Transwell assay: After 24 hours of incubation, USP19-depleted cells were stained for microscopic examination and the number of migratory cells was compared to control cells. The graph shows the number of migratory cells per transwell membrane (n= 4, one-way ANOVA, Dunnett’s multiple comparison test. shRNA#1 p= 0.0187 and shRNA#2 p= 0.0001). (**E**) Wound healing assays: scratching with a pipette tip made a gap on a monolayer of the different cell cultures, and time-lapse imaging monitored the number of migrating cells across the border. After 8 hours, cells exhibited different levels of migration. The graph on the left shows the gap covered area (mm2) after 8 hours (n= 16, one-way ANOVA, Dunnett’s multiple comparison test. shRNA#1 p< 0.0001 and shRNA#2 p< 0.0001) and the images on the right show representative areas in a wound healing experiment at the indicated time points. Scale bar= 100 μm.

### Validation of USP19 as a regulator of cell migration

In order to validate USP19 as a potential regulator of cell migration, we established stable MDAMB231 cell lines transduced individually with two different shRNAs targeting USP19 expression (named shRNA#1 and shRNA#2). Our results showed that both caused a significant reduction in USP19 mRNA and protein levels (Fig. 2B).

It is conceivable that shRNAs promoting cell proliferation may have also been enriched during the functional selection, as they provide cells an advantage during the *in vitro* amplification step. To discard this possibility, we performed proliferation curves of control and USP19-silenced cell lines and calculated doubling rates. We observed no differences between the different cell lines (Fig. 2C, Supp. Fig. 2), providing evidence for a direct role of USP19 in the control of cell migration.

Next, to confirm the effect of USP19 depletion on cell motility, we used two independent assays: transwell migration and wound-healing assays. As shown in Figures 2D and 2E, USP19 knockdown significantly decreased the migratory potential of cells relative to the control cell line. A more detailed analysis on the wound healing assay indicated that wound-edge cells speed and total displacement was significantly reduced in USP19 knockdown cells, and they presented a minor increase in persistence relative to control cells (Supp. Fig. 3). We also compared the effect on migration of USP19 silencing with USP10 silencing, one of the already published candidate genes obtained from our screen [38–40]. Our results indicate that knock down of both genes impair migration to a similar extent (Supp. Fig. 4).

**Figure 3.**
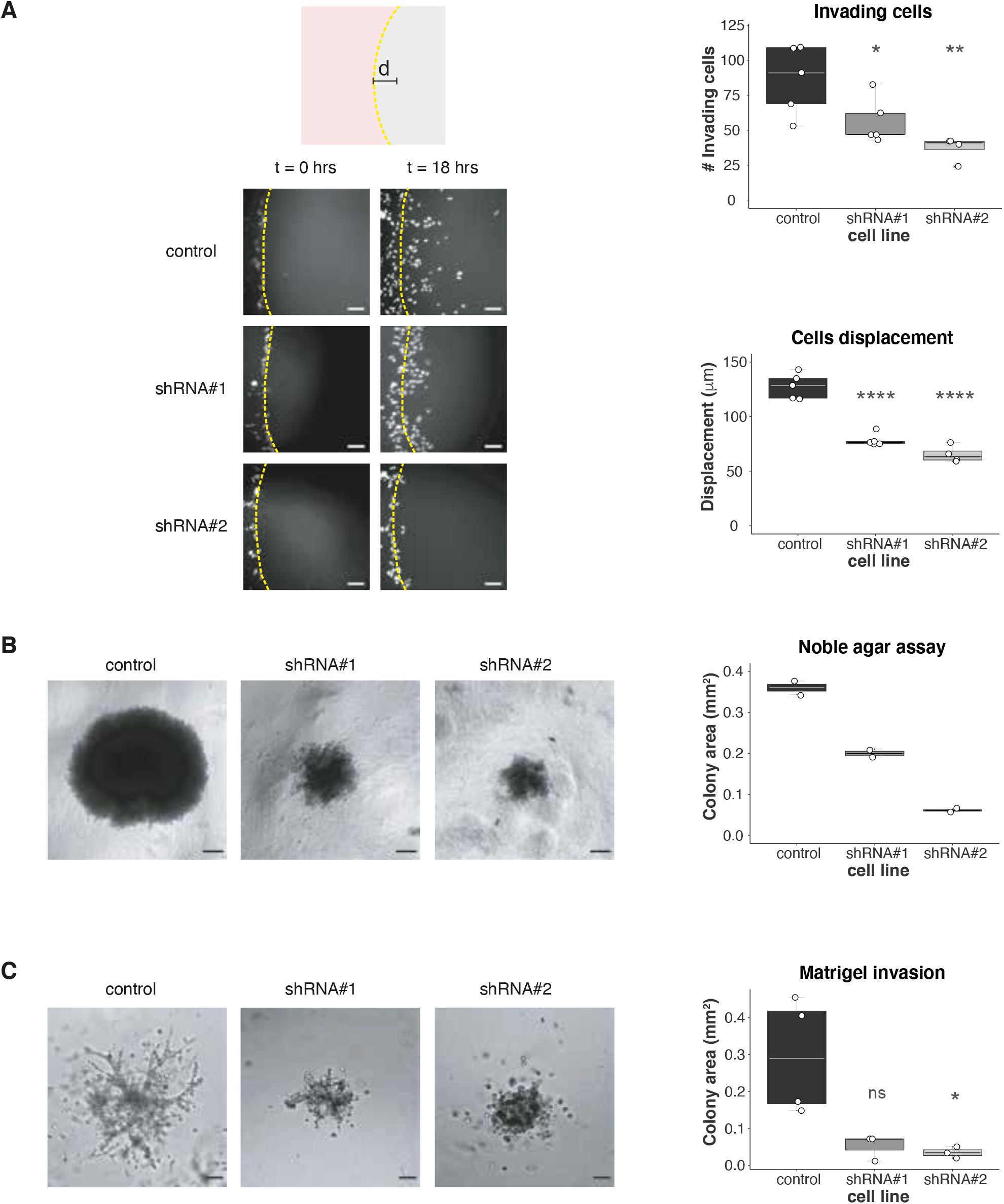
USP19 knockdown reduces cellular invasion. (**A**) Top left: Diagram of agar spot assay. MDAMB231 cells were seeded in wells (pink surface) with drops of solidified agar (gray sphere) and allowed to invade along the bottom surface under the agar. Pictures were taken along the edge (the edge is indicated by a dotted line); the displacement (d) is the extent of invasion under agar from the spot edge until the end of the experiment. Bottom Left: Representative area showing cell invasion into an agar spot at the indicate time points. Top right: Quantification of the mean number of invading cells per spot (n≥ 4, one-way ANOVA, Dunnett’s multiple comparison test. shRNA#1 p= 0.0497 and shRNA#2 p= 0.0042) and bottom right: cells mean displacement after 18 hours (n≥ 4, one-way ANOVA, Dunnett’s multiple comparison test. shRNA#1 p< 0.0001 and shRNA#2 p< 0.0001). Scale bar= 100 μm. (**B**) Noble agar assay was used to study 3D culture proliferation and invasion. Left: Representative brightfield images obtained at 6 weeks in culture are shown. Scale bar= 150 μm. Right: Colony size was calculated at the end of the experiment (n= 2). (**C**) Left: Representative area showing cell invasion in a Matrigel 3D experiment after 5 days in culture. Right: Colony area was calculated at the end of the experiment (n≥ 3, Kruskal-Wallis, Dunn’s multiple comparison test. shRNA#1 p= 0.1033 and shRNA#2 p= 0.0348). Scale bar= 150 μm.

Altogether, these experiments indicate that USP19 silencing affects cell migration *in vitro*. We further confirmed our findings using another highly invasive breast cancer cell line (Supp. Fig. 5).

### USP19 knockdown impairs invasion

Cell motility is often associated with increased tumor cell invasion and is a characteristic trait of aggressive tumor cells [41, 42]. Based on the observation that modulation of USP19 expression regulates migration, we decided to investigate the effect of USP19 depletion on tumor cell invasion.

To this end, we used three different experimental approaches. We first analyzed the ability of cells to invade agar spots. Our results show that USP19 knockdown significantly reduced the number of invading cells as well as their total displacement, compared to the control cell line (Fig. 3A).

We next performed a 3D growth assay by seeding cells at low confluence into noble agar, an anchorage-independent matrix. After 6 weeks in culture, the control cell line formed bigger colonies compared to USP19-silenced cell lines (Fig. 3B), indicating that colonization, matrix invasion and anchorage independent growth in these conditions is partially impaired in cells where USP19 expression is reduced.

Finally, to further characterize USP19 depletion on tumor cell invasion, we assessed growth and invasion into a reconstituted extracellular matrix that provides anchorage (Matrigel^®^). Both cell lines expressing USP19 shRNAs showed colonies with a significantly smaller size than the control cell line (Fig. 3C), indicating that USP19 is required for efficient invasion even when anchorage is provided.

In a similar way to the migration experiments, we further validated our results using another breast cancer cell line (Supp. Fig. 5).

Collectively, our results indicate that USP19 knockdown inhibits tumor cell invasion *in vitro*.

### USP19 overexpression enhances migration and invasion

In order to complement our analysis on the putative role of USP19 as a positive regulator of migration and invasion, we analyzed the effect of USP19 overexpression in a poorly migratory and non-invasive breast cancer cell line (MCF7).

For this purpose, we stably transfected MCF7 cells with a USP19 overexpressing plasmid (Fig. 4A), and then performed wound healing assays. As shown in Figure 4B, USP19 overexpression induced a significant increase in the gap covered area, compared to the control cell line.

**Figure 4.**
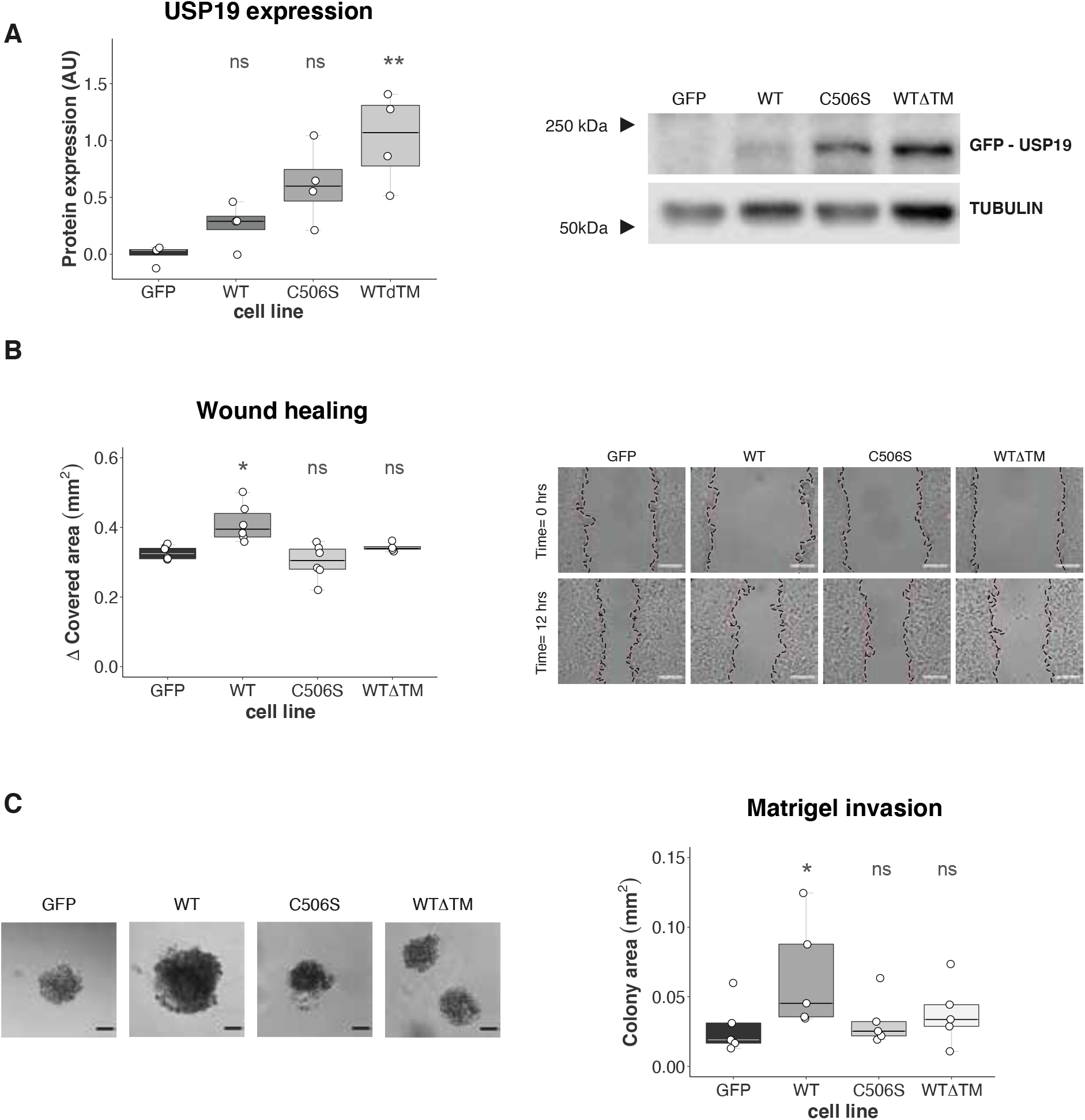
USP19 wild Type overexpression enhances migration and invasion in MCF7 cells. (**A**) Different constructs of USP19 were overexpressed in MCF7 cells, and their expression confirmed by Western Blotting (n=4, Kruskal-Wallis, Dunn’s multiple comparison test. WT p>0.9999, C506S p= 0.0777 and WTΔTM p= 0.0070). (**B**) Migratory potential was evaluated by wound healing assay. Left: gap covered area (mm2) after 12 hours (n≥ 4, Kruskal-Wallis, Dunn’s multiple comparison test. WT p= 0.0107, C506S p>0.9999, WTΔTM p>0.9999) and right: representative areas in a wound healing experiment at the indicated time points. Scale bar= 100 μm. (**C**) Matrigel invasion was assessed over a 30 days period. Left: Representative brightfield images obtained at the end of the experiment using a 10X objective are shown; scale bar= 100 μm. Right: colony size was calculated (n≥ 4, Kruskal-Wallis, Dunn’s multiple comparison test. WT p= 0.0485, C506S p>0.9999, WTΔTM p>0.9999).

As a control, we overexpressed a catalytically mutant version of USP19 [28, 43–47] and a mutant lacking USP19 transmembrane domain (Supp. Fig. 6). In contrast to USP19 wild type, we did not detect any substantial increase in migration in either of these mutants compared to the control cell line (Figure 4B).

This result further supports the hypothesis that USP19 is a positive regulator of migration, and it provides evidence that this phenotype is dependent on its catalytic activity and on its subcellular localization.

Next, we analyzed the effect of USP19 overexpression on invasion and growth into a reconstituted extracellular matrix (Matrigel^®^), using similar experimental settings as described before. We observed a significant increase in colony areas when comparing wild type USP19 overexpressing cells to the control cell line (Fig. 4C). In accordance with our previous results, the USP19-dependent increase in invasion is also determined by its catalytic activity and presence of the transmembrane domain (Fig. 4C).

### USP19 regulates invasion *in vivo*

To further characterize USP19-dependent control of cell invasion *in vivo*, we performed subcutaneous orthotopic xenotransplants and experimental metastasis assays in immunocompromised mice (NOD/SCID).

First, we injected MDAMB231 control or USP19-silenced cells subcutaneously in the mammary fat pad of female mice and monitored tumor growth every 2-3 days. Tumor growth curves analysis indicated that those generated from control cells were significantly more volumetric than the ones originated from USP19-silenced cells (Fig. 5A, left and Supp. Fig. 7). Moreover, Kaplan-Meier curves for tumor-free survival indicated that cells expressing either of the shRNAs targeting USP19 generated fewer tumors compared to the control cell line (Fig. 5A, right and Supp. Table 2). In addition, we observed similar results using another breast cancer cell line (Supp. Fig. 8 and Supp. Table 2).

**Figure 5.**
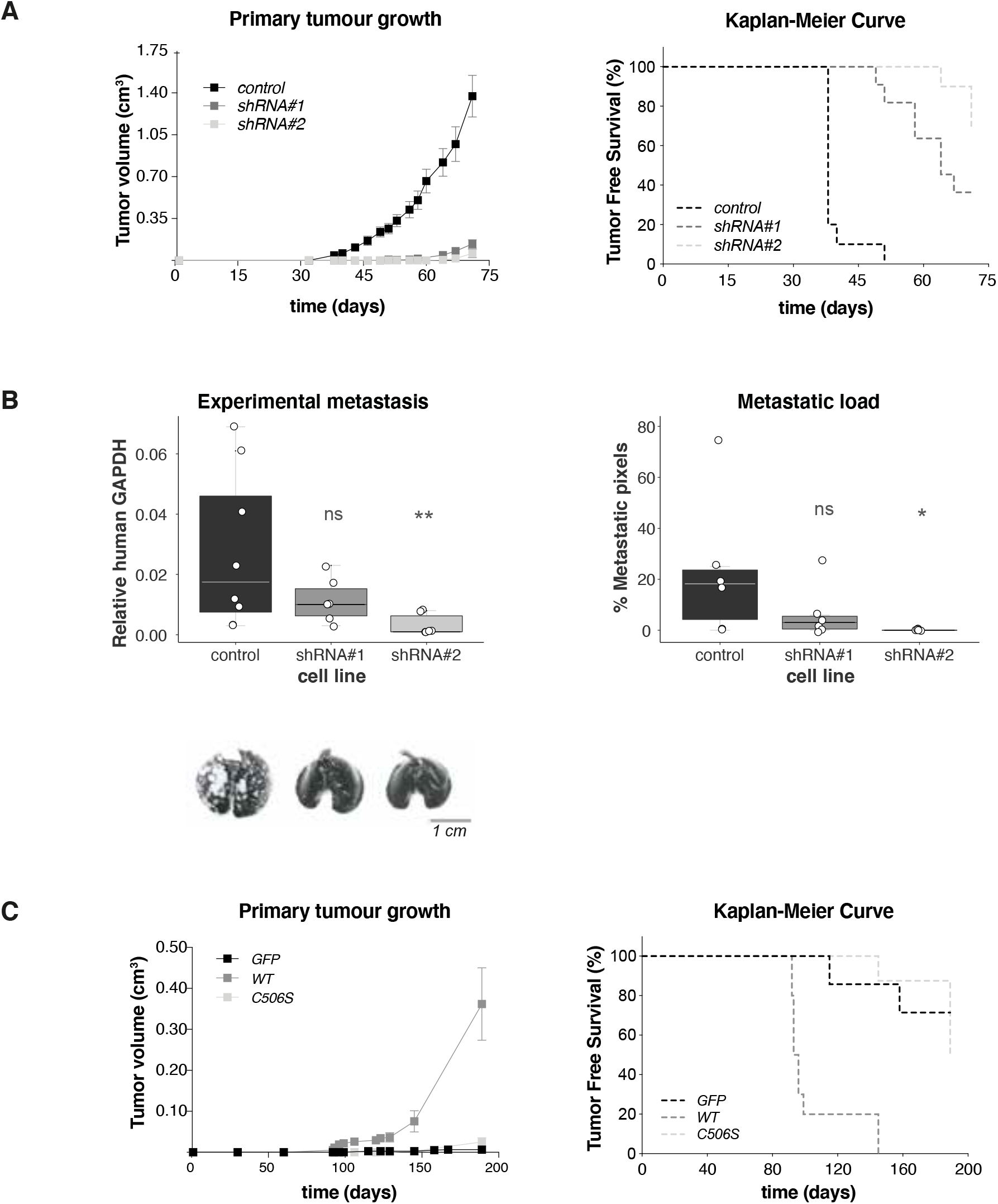
Analysis of USP19 expression relevance using mice models. (**A**) Downregulation of USP19 attenuates tumorigenicity in vivo: Control or USP19-silenced MDAMB231 cells were subcutaneously inoculated into the mammary fat pads of female NOD/SCID mice and tumor growth monitored every 2-3 days. Left: tumor volume was calculated at the indicated time points (results show mean value ± S.E.); right: Kaplan-Meier curves were built for Tumor Free Survival (TFS) over time (n≥ 10, Log-Rank (Mantel-Cox) test, shRNA#1 p< 0.0001 and shRNA#2 p< 0.0001). (**B**) Silencing effects of USP19 on experimental metastasis assays: NOD/SCID male mice were inoculated with MDAMB231 USP19-silenced cells through tail vein injection and after 2 months, lungs were harvested. Top left: metastatic foci were estimated by qPCR human DNA quantification (n≥ 6, Kruskal-Wallis, Dunn’s multiple comparison test. shRNA#1 p> 0.9999 and shRNA#2 p= 0.0032). Bottom left: representative lung images stained with Indian ink at the end of the experiment are shown. Scale bar= 1 cm. Right: metastatic load quantification was performed by evaluating lung Hematoxylin & Eosin stained slides. We used a lesion-based analysis of percent of metastatic pixels to compare the differences in metastatic load produced on the lungs by the MDAMB231 cell lines (n= 6, Kruskal-Wallis, Dunn’s multiple comparison test. shRNA#1 p> 0.9999 and shRNA#2 p= 0.0299). (**C**) USP19 catalytic activity is needed for tumorigenicity in vivo: control, WT or C506S mutant versions of USP19 overexpressing MCF7 cells were subcutaneously inoculated into the mammary fat pads of female NOD/SCID mice. Left: tumor volume was calculated at the indicated time points (results show mean value ± S.E.); right: Kaplan-Meier curves were built for Tumor Free Survival (TFS) over time (n≥ 7, Log-Rank (Mantel-Cox) test, WT p< 0.0001 and C506S p= 0.5307).

Second, we analyzed USP19’s role in the regulation of tumor cell lung colonization. For that purpose, we inoculated control or USP19-silenced MDAMB231 cells through tail vein injection and harvested the lungs two months later. As shown in Figure 5B, USP19 depletion inhibits tumor foci formation *in vivo*, as evaluated by human DNA quantification (left) and metastatic load quantification in Hematoxylin & Eosin stained lung sections (right, and Supp. Fig. 9). We observed the same trend when another breast cancer cell line was used (Supp. Fig. 8).

Last, we repeated the same type of tests using MCF7 cells in similar experimental conditions. We subcutaneously injected control cells or cells expressing either wild type or catalytically mutant versions of USP19 in female mice.

In agreement with our *in vitro* experiments, wild type USP19-expressing cells formed tumors in all injected mice, whereas mice injected with cells expressing the catalytic mutant did not show signs of tumor growth (Fig. 5C, Supp. Fig. 7 and Supp. Table 2). Since these cell lines showed no difference in proliferation rates in two dimensions (Supp. Fig. 10) and the fact that the MCF7 cell line does not usually form tumors unless an external estrogen source is supplied, this result highlights the importance of USP19 for tumor development and onset.

Altogether, we concluded that USP19 is important for *in vivo* colonization and tumor growth. In addition, our results indicate that USP19 catalytic activity and transmembrane domain are required for its stimulatory effect on cell motility.

### USP19 regulates LRP6 protein levels in breast cancer cells

In order to study the putative mechanism of action responsible for USP19 migration and invasion regulation, we performed an *in silico* analysis on breast cancer mRNA expression using publicly available datasets. Our results revealed that high USP19 expression levels correlate with the activation of the Wnt pathway (Fig. 6A, B and C), among others. This result was in concordance with previous observations by Perrody and collaborators, which demonstrated that USP19 stabilizes LRP6, a Wnt pathway coreceptor, and that this interaction affected downstream Wnt signaling capacity [32].

**Figure 6.**
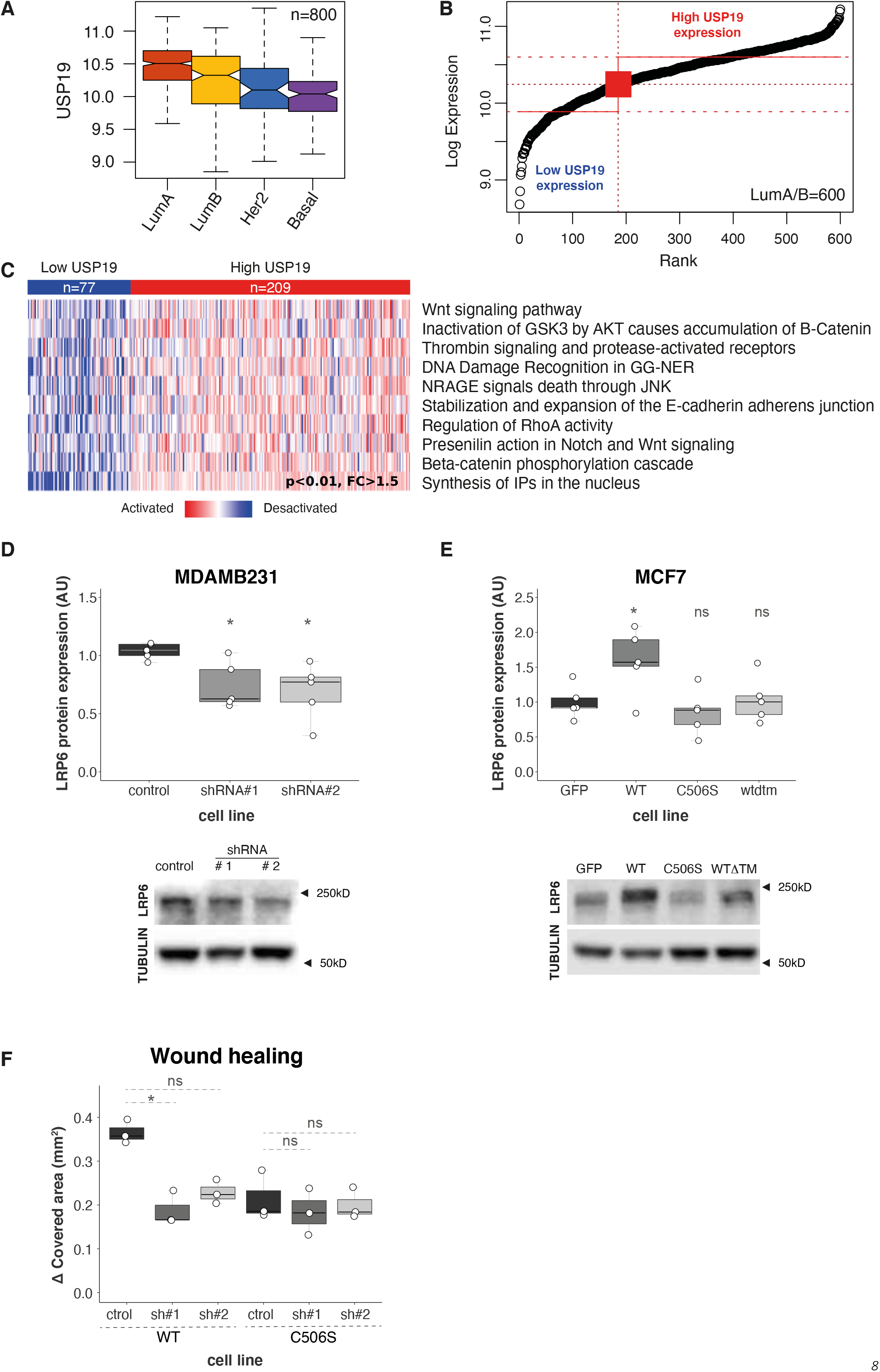
USP19 mechanism of action. An in silico study was performed in order to analyze the relationship between USP19 expression levels and different pathway activation status. (**A**) USP19 mRNA expression among primary breast carcinomas according to their intrinsic subtype. Expression analysis showed a consistent up-regulation in luminal A and B subtypes compared with basal-like and Her2 subtypes. (**B**) Luminal A/B primary breast cancers divided into low (n= 77) or high (n= 209) USP19 mRNA expression levels. (**C**) Significantly activated pathways among Luminal A/B tumors with high USP19 mRNA expression (n≥ 77, SAM test, p< 0.01). Western blotting was performed in order to analyze LRP6 protein expression in breast cancer cells upon USP19 genetic manipulation. (**D**) Top: Western Blot quantification in control or USP19 silenced MDAMB231 cells (n=5, one-way ANOVA, Dunnett’s multiple comparison test. shRNA#1 p= 0.0492 and shRNA#2 p= 0.0226), bottom: representative image of a blot. (**E**) Top: Western Blot quantification in MCF7 cells overexpressing control or GFP-tagged USP19 constructs (n=5, one-way ANOVA, Dunnett’s multiple comparison test. WT p= 0.0484, C506S p= 0.8469 and WTΔTM p= 0.9968), bottom: representative image of a blot. (**F**) Wound healing assays were performed in order to analyze endogenous LRP6 silencing effects in MCF7 cells overexpressing WT or C506S mutant versions of USP19. Cells were stably transduced with control vector (‘ctrol’, PLKO.1 empty vector), or shRNAs targeting LRP6 (sh#1 and sh#2). Scratching with a pipette tip made a gap on a monolayer of the different cell cultures, and time-lapse imaging monitored the number of migrating cells across the border. The graph shows the gap covered area (mm^2^) after 8 hours (n=3, Kruskal-Wallis and Dunn’s multiple comparison test for WT or C506S overexpressing MCF7 cell lines, analyzed separately. WT overexpressing MCF7 cell line: sh#1 p= 0.0341 and sh#2 p= 0.2021. C506S overexpressing MCF7 cell line: sh#1 p= 0.7422 and sh#2 p> 0.9999).

Based on these results, we analyzed LRP6 protein steady state levels upon USP19 genetic silencing or overexpression. In accordance with Perrody *et al.* [32], our results indicate that LRP6 protein levels decrease upon USP19 silencing in MDAMB231 (Figure 6D) and increase in wild type USP19-overexpressing MCF7 cells, but not in cells expressing catalytically dead or cytoplasmic mutant versions (Fig. 6E). This correlation was also observed when using another breast cancer cell line (Supp. Fig 11).

In order to test the functional relation between USP19 and LRP6, we then analyzed the effect of LRP6 endogenous silencing in MCF7 cells overexpressing USP19. Our results indicated that wild type USP19-induced increase in migration was reverted by LRP6 shRNAs stable expression (Fig. 6F).

Altogether, our results indicate that the axis USP19/LRP6, rather than the absolute level of expression of USP19 (Supp. Fig. 11), is key to regulate the migratory potential of breast cancer cells.

### Survival analysis of USP19 expression in early breast cancer patients

Finally, we analyzed USP19 protein expression in a cohort study of early breast cancer patients (T1-2, N0, M0; n= 168) with long-term follow-up. Kaplan-Meier plots showed that overexpression of USP19 was associated with a significantly lower frequency of distant relapse free survival (DRFS), while no significant correlation with disease free survival (DFS) was observed (Fig. 7A and B).

**Figure 7.**
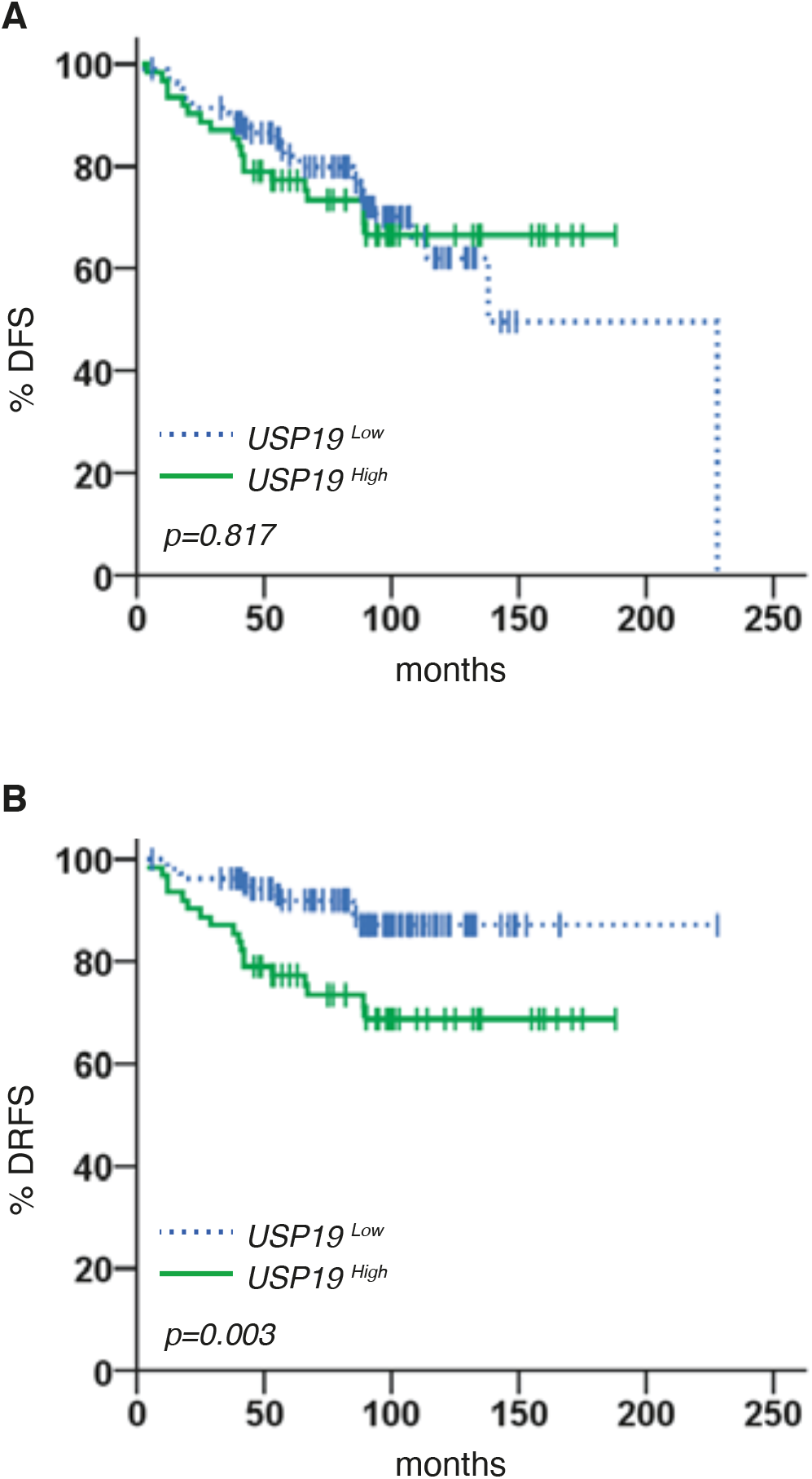
Prognostic value of USP19 protein expression in breast cancer patients. Kaplan-Meier estimates of DFS and DRFS in patients with early breast cancer tumors, according to high (solid green lines) and low (dashed blue lines) expression of USP19. (**A**) 19 out of 62 patients (30.6%) harboring USP19^High^ tumors and 30 out of 106 patients (28.3%) with USP19^Low^ tumors had a disease relapse (n= 168, Log-Rank (Mantel-Cox) test, p= 0.817). (**B**) Distant metastases developed in 18 out of 62 (29.0%) and 11 out of 106 (10.4%) of patients with USP19^High^ and USP19^Low^ tumors, respectively (n= 168, Log-Rank (Mantel-Cox) test, p= 0.003).

Multivariate analysis of DRFS, adjusted for other prognostic factors, revealed that USP19^High^ was an independent prognostic predictor of DRFS (Table 1).

**Table 1.** USP19^High^ is an independent prognostic predictor of DRFS. Multivariate analysis of USP19 expression in breast tumors indicated that USP19^High^ is an independent prognostic predictor of DRFS. HR: Hazard Ratio, CI: Confidence Interval (n= 168, Cox’s proportional hazard model, DFS p= 0.926, DRFS p= 0.032).

| Variable | HR | 95% CI | P* |
| --- | --- | --- | --- |
| **Disease-free survival** |  |  |  |
| Tumor size, cm (>2 vs ≤ 2) | 1.0 | 0.6-1.9 | 0.888 |
| Tumor grade (1 vs 2-3) | 1.1 | 0.5-2.8 | 0.793 |
| ER (negative vs positive) | 1.6 | 0.6-4.2 | 0.302 |
| PR (negative vs positive) | 1.1 | 0.5-2.6 | 0.853 |
| Ki-67 (positive vs negative) | 1.1 | 0.6-2.0 | 0.818 |
| HER-2 (positive vs negative) | 1.9 | 0.9-4.2 | 0.112 |
| USP19 (low vs high) | 1.0 | 0.6-1.9 | 0.926 |
| **Distant Relapse-Free Survival** |  |  |  |
| Tumor size, cm (≤ 2 vs >2) | 1.4 | 0.6-3.1 | 0.413 |
| Tumor grade (2-3 vs 1) | 3.6 | 0.5-27.0 | 0.221 |
| ER (negative vs positive) | 2.7 | 0.9-8.8 | 0.090 |
| PR (positive vs negative) | 1.8 | 0.6-5.3 | 0.313 |
| Ki-67 (positive vs negative) | 1.1 | 0.5-2.4 | 0.808 |
| HER-2 (positive vs negative) | 1.9 | 0.8-4.6 | 0.176 |
| USP19 (high vs low) | 2.4 | 1.1-5.4 | 0.032* |

Altogether these findings indicate that, in accordance with our *in vitro* and *in vivo* studies, USP19 represents a new predictor of distant metastasis formation in early breast cancer patients.

## DISCUSSION

Migration occurs in a wide variety of physiological conditions, and alterations in its regulation are associated with different pathologies, including cancer [3, 4, 48]. In this disease, mortality is associated primarily with tumor growth at secondary sites, and effective therapies to block the metastatic cascade are lacking [6]. Tumor cells need to migrate, invade and colonize new niches prior to metastasis, making it a vital trait of malignancy. Indeed, a recent work indicated that migration, rather than proliferation, is strongly associated with breast cancer patient survival [49].

Therefore, modulation of genes that regulate migration and invasion could find application for the treatment of cancer. In line with this reasoning, we chose to screen for genes that positively regulate motility within the ubiquitination pathway, as this cascade is currently emerging as an attractive therapeutic target in drug development [50–53]. Here we report the identification of USP19, a deubiquitinating enzyme, as a positive regulator of migration in breast cancer. USP19 was initially characterized as a DUB predominantly localized in the cytosol in association with Hsp90 and other chaperones [31]. USP19 has been associated with the regulation of the half-life of several proteins that participate in different cellular processes [27, 32, 45, 46, 54–63].

Our *in vitro* validation experiments showed that USP19 depletion did not affect cell proliferation in agreement with Lu *et al.* [64], but directly inhibited cellular migration. In addition, we observed that USP19 knockdown impaired invasion, as evaluated by agar drop assays and three-dimensional basement membrane cultures (Figs. 2, 3 and Supp. Fig. 5). When we analyzed growth into noble agar, the size of the resulting colonies was smaller than control cells, indicating that USP19 silencing also reduced anchorage-independent growth (Fig. 3). In all cases, we observed a correlation between the extent of USP19 silencing and the reduction of migration and invasion potential. These results are in agreement with USP19-silencing deleterious effects in growth and development in Zebrafish embryos [65].

To further confirm our results, we analyzed how USP19 overexpression affected migration and invasion, using a poorly migratory cell line. In agreement with our depletion experiments, USP19 overexpression induced an increase in cellular migration, invasion and growth in three-dimensional basement membrane cultures.

These effects were dependent on USP19 its subcellular-localization, and on the presence of a highly conserved cysteine at the catalytic site and mutation of this residue abolished USP19-induced migration and invasion. Taken together, these *in vitro* results suggest that USP19 expression levels are associated with the regulation of motility, invasion, and anchorage-independent growth in breast cancer cell lines.

Our *in vivo* studies using immunocompromised mice demonstrated that USP19 silencing decreased cell engraftment and tumor growth, as well as colonization into the lungs (Fig. 5A and Supp. Fig. 8B). On the contrary, overexpression of wild type USP19, but not its catalytically deficient mutant version, promoted tumor growth (Fig. 5B). This is compatible with the requirement of USP19 catalytic activity for local invasion and growth in three dimensions, both *in vitro* and *in vivo*. In line with these results, we observed a marked increase in USP19 mRNA expression in cells growing in tumors compared to the same cells in culture dishes (Supp. Fig 12). These results are compatible with a requirement for higher levels of USP19 to support three-dimensional invasion and growth, highlighting the possible existence of a specific regulation of USP19 in a context where cells need to invade.

Finally, a retrospective study conducted on human breast tumor samples indicated that high USP19 protein levels are associated with high-risk for metastatic relapse in patients diagnosed with early breast cancer (Fig. 7).

Altogether these results provide evidence indicating that USP19 has great potential as a therapeutic target for drug development in breast cancer treatment.

In this regard, there is considerable scientific evidence demonstrating that DUBs exhibit strong substrate selectivity, which can be advantageous to ensure high efficacy and low adverse effects. Moreover, the design and development of a selective enzyme inhibitor is easier than generating an enzyme activator due to competitive inhibition and modeling of substrates [66]. In fact, numerous inhibitors for DUB activities have been already identified [67–71] and an orally bioavailable compound that inhibits USP19 activity has recently been developed [72].

USP19 was previously related to proliferation and growth in Ewing’s sarcoma cells, a specific type of cancer characterized by a reciprocal translocation and fusion of the EWSR1 and the FLI1 genes. The authors demonstrated that USP19 specifically stabilized EWS-FLI1 fusion oncoprotein, but not EWSR1 or FLI1 proteins [45], therefore indicating that the specific molecular target of USP19 in this context is specific for this type of cancer and the results cannot be easily extrapolated to other contexts.

To our knowledge, USP19 molecular mechanism of action in the regulation of migration and invasion in breast cancer cells was not investigated before.

Our results demonstrated that USP19 expression correlates with tumor growth and invasion. Supporting this, we analyzed e-cadherin protein expression levels in the samples of our retrospective study and observed an inverse correlation between USP19 and e-cadherin expression (n= 168, Spearman correlation analysis, rho= −0.180, p= 0.032, Supp. Table 3). In agreement with our results, previous reports demonstrated that low e-cadherin expression holds a prognostic value as a predictor of poorer prognosis and more aggressive phenotypes in breast cancer [73, 74].

Moreover, we performed an *in silico* analysis on breast cancer mRNA expression publicly available datasets, which revealed that high USP19 expression levels correlate with the activation of the Wnt pathway (Fig. 6 A, B and C). This is consistent with a recent work which showed that USP19 regulates LRP6 stability, a co-receptor of the Wnt signaling cascade [32]. Particularly in breast cancer, LRP6 is overexpressed in around a third of the patient samples, and its overexpression has been proposed as a distinctive feature of a specific class of breast cancer subtype [75].

In this regard, our experiments show that LRP6 expression positively correlates with USP19 protein levels in breast cancer cells (Fig. 6D and E, and Supp. Fig. 11) and that overexpression of a catalytically dead mutant or a cytoplasmic version of USP19 has no effect on LRP6 (Fig. 6E), in concordance with previous results [32]. Moreover, this molecular mechanism is specific associated with USP19 modulation and it is not a general effect as a result of change in migration, since downregulation of USP10 and its concomitant reduction in migration does not alter LRP6 protein levels (Supp. Fig. 4C). In all, our results are compatible with former experiments that demonstrated that LRP6 downregulation in breast cancer cell lines reduces their migratory and invasive potential [76], as well as their ability to form colonies in soft agar [75]. More importantly, we show that endogenous LRP6 silencing abolishes USP19 overexpression-induced increase in migration (Fig. 6F). Consequently, our results indicate that the functional interaction between USP19 and LRP6 is key for the regulatory effect that USP19 exerts on the modulation of breast cancer cells migration and invasion.

Opposite to our findings, Hu and collaborators very recently demonstrated that USP19 negatively regulates proliferation and migration in clear cell renal carcinoma [77]. In this type of cancer, the most relevant USP19 isoform is uc003cvz.3, which is mainly localized in the cytoplasm [78]. Based on our data showing that the control of cell migration in breast cancer cells is mainly exerted by the transmembrane USP19 isoform, it is plausible to assume that this difference could contribute to explain the divergent role that USP19 plays in these two different cellular contexts.

For all the reasons expressed before, we conclude that USP19 is relevant for the regulation of breast cancer cell dissemination and its expression levels correlate with high risk of metastases development, and could therefore represent a novel target for the management of breast cancer metastatic disease, in particular when LRP6 expression is relevant for determining patients’ outcome.

## MATERIALS AND METHODS

### Cell lines and cell culture

The human breast cancer cell lines MCF7, MDAMB231 and MDAMB436 and the Hek293T cells were obtained from the ATCC and cultured in Dulbecco’s modified Eagle’s medium (DMEM) (Gibco) supplemented with 10% fetal bovine serum (FBS) (Natocor, Córdoba, Argentina), 50U/ml penicillin-streptomycin and 200 μM L-glutamine at 37°C and 5% CO_2_ in a humidified incubator. ATCC uses morphology, karyotyping, and PCR based approaches to confirm the identity of human cell lines. Mycoplasm contamination was evaluated monthly by PCR, and cell lines were cultured less than three months.

### shRNA screening and plasmid transfections

A pool of plasmids encoding 1,885 shRNAs targeting 407 different genes related to the ubiquitination pathway (UB/DUB library) in the pLKO.1 backbone produced by The RNAi Consortium (TRC, Sigma-Aldrich, St. Louis, MO) were obtained from the University of Colorado Cancer Center Functional Genomics Shared Resource. 1 mg of the shRNA library plasmid DNA at 100 ng/mL was mixed with 4 mg of packaging plasmid mix (pD8.9 and pCMV-VSVG lentiviral packaging plasmids at a 1:1 ratio) and incubated with 30 mg of polyethylenimine for 15 min at RT. The entire mixture was then added to a 100 mm dish containing Hek293T packaging cells at 75% confluence. 6 hours after transfection, media on cells was replaced with complete DMEM and 48 hours after media replacement, the supernatant from each dish of packaging cells (now containing lentiviral library particles) was filtered through 0.45 mm cellulose acetate filters and stored at −80°C until use. Before performing the screen, MOI determination of the lentivirus stock was carried out using different dilutions. Target cells were seeded at 8×10^4^ cells/well in 6-well plates and then transduced with the lentivirus. After 48 hours the infective media was removed, and target cells were selected for 5 days with a 0.5 μg/ml puromycin DMEM medium. The amount of virus required to maintain 10% survival was used. These infection conditions were essential for each cell to be transduced with less than one lentivirus particle expressing a single shRNA. Target cells were transduced with a 1:27 dilution of the lentivirus stock and were then combined at the time of harvest to reach a starting number of 4,4×10^6^ cells per condition (~300X coverage of the library complexity). 48 hours after transduction, the media was replaced with puromycin selection media and cells were then propagated for 14 days before use, in order to select out those shRNA targeting essential genes.

For single shRNA transduction, TRCN0000051715 and TRCN0000051716 (USP19 shRNA# 1 and 2, respectively), TRCN0000033406 and TRCN0000033408 (LRP6 shRNA# 1 and 2, respectively), and SHC001 (control) PLKO.1 vectors were used (obtained from the University of Colorado Cancer Center Functional Genomics Shared Resource); no MOI determination was performed.

For overexpression experiments, transfections were performed using Lipofectamine 2000 reagent (Invitrogen, Carlsbad, CA) according to the manufacturer’s protocol. GFP tagged wild type and catalytically dead mutant (C506S) USP19 plasmids were a kind gift of Dr. Sylvie Urbé (University of Liverpool, UK), and GFP tagged ΔTM USP19 plasmid was obtained by generating a premature stop by mutagenesis PCR from wild type USP19 vector.

### Mutagenesis

Mutagenesis PCR amplification was performed with the KAPA HiFi System (Kapa Biosystems) following the manufacturer’s instructions. 1 μL DpnI enzyme (20 U, New England Biolabs #R0176L) was added to the 25 μL PCR mixture immediately after the final extension and incubated at 37°C for 1 hour. The digestion product was then purified with the QIAquick PCR Purification Kit (Qiagen) following the manufacturer’s instructions, and 5 μL of the mixture was used to transfect competent E coli DH5α. Mutations that generated a stop codon at aminoacid 1290 were corroborated by sequencing and by checking expression in U2OS cells under a fluorescence microscopy (Supp. Fig 6). Mutagenesis primers sequences are as follows: 5’-GATGAGGGCTGCCTCCGGTAGTAGTAGCTGGGCACCGTGGCGG-3’, 5’-CCGCCACGGTGCCCAGCTACTACTACCGGAGGCAGCCCTCATC-3’.

### Transwell migration assay

All cell lines were starved during 24 hours in assay medium (growth medium containing 0.1% FBS). The starved cells were trypsinized, 5×10^4^ cells were added to the top chamber of 24-well transwells (8 μm pore size membrane; BD Bioscience, Bedford, MA), and assay medium was added to the bottom chambers and incubated for 24 hours (10% FBS). After non-migratory cells removal, membranes were fixed, stained with 4ʹ,6-diamidino-2-phenylindole, and mounted. The whole membrane was then imaged using a Zeiss Axio Observer Z1 Inverted Epi-fluorescence microscope with montage function. Image analysis was performed with Fiji software, using an automated analysis macro to measure the number of nuclei per transwell.

For the migration screen, 6.6×10^5^ starved MDAMB231 cells expressing the shRNA library (or control cell line) were plated onto 6-well transwell chambers (8 μm pore size membrane; BD Bioscience, Bedford, MA) in assay medium. After a 24-hour incubation, the non-migratory cells were collected from the upper chamber, propagated and allowed to re-migrate eleven times for enrichment purposes (the non-migratory cells of each migration experiment were used for the subsequent cycle of enrichment). The non-migratory cells of 8 transwells were combined per each selection cycle to ensure a > 700 library coverage. Simultaneously, the percentage of non-migratory cells in each cycle was determined in 24-well plates as described before, using 8 μm pore size membrane transwells (BD Bioscience, Bedford, MA).

### shRNA library preparation and sequencing

The library preparation strategy uses genomic DNA and two rounds of PCR in order to isolate the shRNA cassette and prepare a single strand of the hairpin for sequencing by means of an XhoI restriction digest in the stem-loop region.

We used barcoded half-hairpin sequences for the identification of shRNAs from every enrichment cycle. The procedure was performed as described previously [79]. After purity analysis of the sequencing library, barcode adaptors were linked to each sample to allow a multiplexing strategy. A HiSEQ 2500 HT Mode V4 Chemistry Illumina instrument was used for that purpose and each sample was quantified and mixed together at a final concentration of 10 ng/mL. Samples were sequenced with a simple 1×50 run and on average 1.2×10^6^ reads were obtained per sample (> 600X shRNA library complexity).

### shRNA screen analysis

shRNA data were analyzed in a similar fashion to RNA-seq data. Briefly, quality control was performed with FastQC, reads were trimmed to include only shRNA sequences using FASTQ trimmer and filtered with the FASTQ Quality Filter. Reads were then aligned to a custom reference library of shRNA sequences using TopHat2.

### Quantitative PCR

Total RNA was extracted from cell lines or tumor tissues using TRIzol reagent (Invitrogen, Thermo Fisher Scientific) according to the manufacturer’s instructions and complementary DNA synthesis was carried out using M-MLV reverse transcriptase in the presence of RNasin RNase inhibitor (Promega) and an oligo(dT) primer (Invitrogen).

Total DNA was extracted from cell lines or fixed tissues using Qiagen DNeasy Blood and Tissue kit with the optional RNAse A treatment step and following the manufacturer’s instructions.

Quantitative real-time PCR amplification, using specific primer sets at a final concentration of 300 nM, was carried out using the FastStart Essential DNA Green Master kit (Roche) at an annealing temperature of 60°C for 35 cycles, and a CFX96 PCR Detection System (Biorad). Expression was calculated for each gene by the comparative CT (ΔCT) method with GAPDH for normalization.

Sequences and expected product sizes are as follows: Usp19 sense 5ʹ-CAAATGTTCTCATCGTGCAGCTC-3ʹ, antisense 5ʹ-CTTGCTCAGGTCCAGGTTCCTAACA -3ʹ (110 bp) and GAPDH sense 5’-TGCACCACCAACTGCTTAGC-3ʹ, antisense 5ʹ-GGCATGGACTGTGGTCATGAG-3ʹ (87 bp) for mRNA expression analysis. (PCR primers are all intron spanning).

Human GAPDH sense 5ʹ-TACTAGCGGTTTTACGGGCG-3ʹ, antisense 5ʹ-TCGAACAGGAGGAGCAGAGAGCGA-3ʹ (166 bp) and mouse GAPDH sense 5’-CCTGGCGATGGCTCGCACTT -3ʹ, antisense 5ʹ-ATGCCACCGACCCCGAGGAA -3ʹ (232 bp) for DNA expression analysis.

### Western blot analysis

Cells were lysed in lysis buffer (50 mM Tris-HCl pH 7.4, 250 mM NaCl, 25 mM NaF, 2 mM EDTA, 0.1% Triton-X, with protease inhibitors mix (Complete ULTRA, Roche), 1 mM 1,4-DTT, 1 μM NaOV, 10 nM okadeic acid). Protein concentrations in cell lysates were determined using the BCA assay Kit (Pierce) and then prepared for loading in Laemmli buffer 4X. Equal amounts of protein were separated by 8-12% SDS-PAGE and transferred to PVDF membranes (Millipore-Merck). Membranes were incubated with primary antibodies: rabbit anti-USP19 antibody (Bethyl Cat# A301-587A, 1:2000 dilution), mouse anti-tubulin antibody (Santa Cruz Biotechnology, Cat# sc-398103, 1:3000 dilution), mouse anti-β-actin antibody (Cell Signaling, Cat# 3700, 1:5000 dilution), mouse anti-GFP antibody (Santa Cruz Biotechnology, Cat# sc-9996 B2, 1:1000 dilution) and rabbit anti-LRP6 antibody (Cell Signaling Cat#2560 C5C7, 1:2000 dilution) and HRP-conjugated secondary antibodies: anti-rabbit (GE Healthcare Cat# NA934); anti-mouse (GE Healthcare Cat# NA931, 1:5000 dilution), and then detected using an ECL SuperSignal West Femto y West Pico detection kit (Pierce).

### Crystal violet proliferation assay

1×10^4^ cells were seeded in 96-well plates at time= 0 hours in quadruple. Every 24 hours, medium was removed at each time point, cells were rinsed with 1X PBS and fixed with methanol for 15 minutes. After 4 days, all wells were staining simultaneously with 150 μl of a 0.05% crystal violet solution for 15 minutes. Plates were then rinsed with water, dried out and the remaining crystal violet was solubilized in 150 μl methanol. The absorbance was measured at 595 nm in a plate reader. An exponential growth model for non-lineal regression was used to calculate doubling time.

### Area-based analysis of proliferation rate

This experiment was performed as described previously, with minor modifications [80]. Briefly, 1×10^3^ cells were seeded in 96-well plates and incubated overnight to allow cell attachment. Images were then acquired under bright field illumination every 6 hours for 3 days using a 10X air objective and Zeiss Zen Blue 2011 software for image acquisition. Image analysis was performed with Fiji software, using an automated analysis macro to measure the occupied area by cells. An exponential growth model for non-lineal regression was used to calculate doubling time.

### Wound healing assays

Cells were seeded onto 24-well culture plates at 2,5×10^5^ per well in growth medium. Confluent monolayers were starved during 24 hours in 0.1% FBS supplemented DMEM medium and a single scratch wound was created using a micropipette tip. Cells were washed with PBS to remove cell debris, supplemented with 3% DMEM medium, and incubated at 37°C under 5% CO_2_ to enable migration into wounds. Images were acquired with a Zeiss Axio Observer Z1 Inverted Epi-fluorescence microscope equipped with an AxioCam HRm3digital CCD camera; a Stage Controller XY STEP SMC 2009 scanning stage and an Incubator XLmulti S1 and Heating Unit XL S1 for live imaging incubation. Images were acquired under bright field illumination using a 10X air objective and Zeiss Zen Blue 2011 software for image acquisition. Image analysis was performed with Fiji software, using an automated analysis macro to measure the occupied area by cells, and the results are presented as the wound covered area at the end of the experiment, relative to time= 0 hours. For wound-edge cells analysis, GFP-tagged H2B expressing MDAMB231 cells were used. Images were acquired every 30minutes for 8 hours for three independent experiments, and calculations were performed using cell tracks from each wound edge separately. The number and position of cells were determined using image analysis software, ImageJ/Fiji -‘Trackmate’ plug-in [81], and trajectory analysis was performed using the ‘Chemotaxis and Migration Tool’ plug-in for ImageJ 1.01 (Ibidi).

### Agar invasion assay

The procedure was performed as described previously, with minor modifications [82–84]. A 1% noble agar solution was heated until boiling, swirled to facilitate complete dissolution, and then taken off of the heat. When the temperature cooled to 50°C, 5 μL spots were pipetted onto 96 well cell culture plates and allowed to cool for 20 min at RT under the hood. At this point, 5×10^3^ cells were plated into spot-containing wells in the presence of 10% FBS cell culture media supplemented with 1 μg/ml Hoechst 33258 (Thermo Fisher Scientific) and allowed to adhere for 1 hour. Fluorescent images of the edges of each spot were taken every 20 minutes during 18 hours on an Axio Observer Z1 (Zeiss) Fluorescence Microscope using a 10X magnification air objective, equipped with CCD Axio Cam HRm3 digital Camera, and a XL multi S1 (D) incubation unit plus a XL S1 (D) temperature module to maintain cell culture conditions at 37°C and 5% CO_2_. Acquisition was controlled with Zen Blue 2011 (Zeiss) Software. Graph construction and statistical analysis were performed using GraphPad Prism. The number and position of cells were determined using ImageJ/Fiji -‘Trackmate’ plug-in [81].

### Noble agar assay

This experiment was performed as previously, with minor modifications [85]. A 2 ml mixture of 5,000 cells in assay medium and 0.3% noble agar was seeded onto a 4 ml-solidified bed of 0.6% noble agar on six-well plates. The plates were allowed to solidify and were then incubated at 37°C. The cultures were fed once a week with assay medium. The cultures were imaged, and the number of colonies was counted after 6 weeks.

### Matrigel three-dimensional cell culture

Experiments were carried out based on experimental settings described before [86–89]. Briefly, 1,000 cells were cultured during the length of the experiment in 100 μl basement membrane gels composed of 9.2 μg/μl phenol red-free growth factor reduced Matrigel (BD Bioscience) in 96 well plates. 50 μl of fresh medium were added on top of each gel every three days.

### Mouse tumorigenesis and metastasis models

NOD SCID mice were originally purchased from Jackson Laboratories (Bar Harbor, ME, USA), and bred in our animal facility under a pathogen-free environment. For all experiments, 7/8-week-old mice were used in accordance with protocols approved by the Institutional Board on Animal Research and Care Committee (CICUAL, Experimental Protocol # 63, 22.nov.2016), Facultad de Ciencias Exactas y Naturales (School of Exact and Natural Sciences), University of Buenos Aires.

For *in vivo* mouse tumor studies, 5×10^5^ transduced cells were suspended in 100 μl of sterile 1X PBS and subcutaneously injected in the mammary fat pads of female mice. Tumors were measured every 3 days and tumor volumes were calculated using the following formula: Vol (volume)= ½ (width^2^ * length). Area Under Curve analysis was performed using measurements from mice that were alive at the end of the experiment.

For the experimental metastasis assay, 1×10^6^ cells were suspended in 200 μl of sterile 1X PBS and injected in the lateral tail vein of male mice. Lungs were harvested 60 days post-injection, fixed in buffered formalin and then stored in 70% ethanol until use for DNA quantification (as described before [90]) or paraffin embedding and Hematoxylin and Eosin staining, or insufflated with a 15% India Ink solution and counterstained with Fekete’s solution for macrometastasis exposure and imaging.

### Metastatic load quantification of lung Hematoxylin & Eosin (H&E) stained slides

The presence of tumor nodules was identified by scanning individual lung H&E stained slides with an optical microscope. Digital image files were acquired for each specimen. For the analysis of the lung metastatic area, Adobe Photoshop software was used to determine the percentage of the lung section that was occupied by the tumor. Furthermore, since the cell-surface glycoprotein CD44s is constitutively expressed by MDAMB231 cells, immunohistochemical staining with the anti-human CD44s (HCAM) antibody (clone DF1485; heat induced epitope retrieval in citrate buffer pH 6.0; dilution 1:50, incubation time: 60’) was used to confirm the tumor origin of the lung nodules (Supp. Fig. 9).

### *In silico* analysis of USP19 mRNA expression among the TCGA-BRCA dataset

Pre-processed USP19 expression levels among 800 primary breast carcinomas with intrinsic subtype data and their integrated pathway activities (pathway activity - z score of 1387 constituent PARADIGM pathways) were obtained from the TCGA Breast Cancer (BRCA) dataset at UCSC Xena browser (http://xena.ucsc.edu/). The PARADIGM algorithm integrates pathway, expression and copy number data to infer activation of pathway features within a superimposed pathway network structure extracted from NCI-PID, BioCarta, and Reactome [91].

Briefly, Luminal A/B primary breast cancer group (n= 600) was divided into low (n= 77) or high (n= 209) USP19 expression levels according to the StepMiner one-step algorithm (http://genedesk.ucsd.edu/home/public/StepMiner/).

These two groups were then compared at their integrated pathway activities to identify the most relevant signaling pathways associated with USP19 expression using the SAM test (p< 0.01; Fold Change> 1.5) with MultiExperiment Viewer Software (MeV 4.9).

### Patients and immunohistochemistry

We retrospectively extracted the eligible patients for the study from a consecutive cohort of cases (year range, 1983-2001) diagnosed with primary unilateral breast carcinoma at the Regina Elena National Cancer Institute, Rome, Italy. From the original series, only N0 patients with T1/T2 tumors were included in this study (n= 168). The patient and tumor characteristics can be found in Supplementary Tables 4 and 5. This study was reviewed and approved by the Ethics Committee of the Regina Elena National Cancer Institute, and written informed consent was obtained from all patients. All the patients were treated with quadrantectomy and received radiation therapy, while 90 (53,6%) received chemotherapy associated or not to hormonal therapy, and 58 (34.5%) underwent only hormonal therapy. Patients with HER2-positive tumors did not receive trastuzumab because it was not available during the study period. The median follow-up was 91.5 months (range, 6 - 298 months). Follow-up data were collected from institutional records or referring physicians. During the follow-up, distant relapse was seen in 29 (17.3%) of the patients.

Tissue microarrays (TMA) were constructed by punching 2-mm-diameter cores from invasive breast carcinoma areas, as previously described [92]. TMA sections were incubated overnight with the rabbit anti-USP19 polyclonal antibody (LifeSpan, Cat# LS-C353286, 1:50 dilution) after applying the MW antigen retrieval technique at 750 W for 10 min in 10 mM Sodium Citrate Buffer (pH 6.0). The anti-rabbit EnVision kit (Agilent, CA) was used for signal amplification. For the negative control, the primary antibody was substituted with a rabbit non-immune serum.

The anti-E-cadherin mouse monoclonal antibody (clone HECD-1, 1:50 dilution, 30 min, Zymed Laboratories Inc., San Francisco, CA) was also used. Antigen retrieval was performed as described in the previous paragraph.

The immunohistochemical analysis was carried out by two pathologists (R.L., S.B.) by agreement, with both blinded to the clinicopathological information. The proportion of USP19-positive cells that showed cytosolic positivity was in the range of 4-100%, with a mean ± S.E. of 63.2% ± 3.6. E-cadherin positivity was defined as a membrane-associated, linear pattern of immunoreactivity which decorated the cell membrane entirely.

The immunohistochemical results for the estrogen receptor (ER), progesterone receptor (PR), Ki67, and HER2 status were obtained from the patient hospital records.

### Statistical analysis

Results are presented as Box-and-whisker plots with median interquartile ranges plus minimum to maximum. *n* indicates the number of independent biological replicates. The one-way ANOVA with Dunnett’s multiple-comparison test as well as non-parametric Kruskal-Wallis and Dunn’s Tests were used to compare treatments to their corresponding control, and adjusted p-values are indicated. P-value differences of < 0.05(*), < 0.01(**), < 0.001(***) or < 0.0001(****) were considered statistically significant. GraphPad Prism and SPSS (SPSS version 15.0, Chicago, IL) statistical software were used for the analysis.

The expression of the USP19 protein in patients’ samples was reported as percent of positive cells and dichotomized (high vs. low) according to the ROC analysis. The optimal cut-off parameter for USP19 positive expression was 50%. Consequently, tumors were identified as USP19^High^ (n= 62) with a score above the cut-off threshold, while it was USP19^Low^ (n= 106) with a score below the threshold (Supp. Fig. 13). Pearson’s χ^2^ or Fisher’s exact tests were used to assess the relations between the tumor USP19 protein expression and the patient clinicopathological parameters. Disease-free survival (DFS) was defined as the interval from surgery to the first of the following events: tumor relapse at local or distant sites. Distant relapse-free survival (DRFS) was defined as the time from surgery to the occurrence of distant relapse. The Log-Rank (Mantel-Cox) test was used to analyze differences between the survival curves, and Cox’s proportional hazard model was used to evaluate the association of USP19 expression with survival time, using covariates. The following covariates were computed in the multivariate model: tumor size, tumor grade, and ER, PR, Ki-67, HER2, and USP19 status.

## ACKNOWLEDGEMENTS

This work was supported by grants from the Agencia Nacional de Promoción Científica y Tecnológica (ANPCyT) PICT 2011-2783, PICT2016-2620 and PICT 2018-03688 awarded to MR. JEM was supported by NIH R01CA117907, NIH R01GM120109, NSF MCB-1817582 and NIH P30CA046934 grants, and GS was funded by an AIRC grant: IG2016 id18467. FAR and JHES are postdoctoral fellows of Consejo Nacional de Investigaciones Científicas y Técnicas (CONICET), and EHCR was recipient from a fellowship from Instituto Nacional del Cáncer (INC).

## AUTHOR CONTRIBUTIONS

FAR conceived, designed, and administered the study; acquired, interpreted, and analyzed data, and wrote the manuscript with assistance from JHES and MR. JHES and EHCR were responsible for acquisition, analysis, and interpretation of data. MJ prepared the shRNA library. AP processed screen sequencing data. MAC performed *in silico* pathway activation analysis. BD and GS performed mouse tissue staining, measurements, and data analysis. SB, VDL, GS and RL performed patients’ biopsies staining, data collection and analysis. JEM provided resources, helped in funding acquisition and coordinated the NGS sequencing and analysis processes. MR conceptualized, designed, supervised, and administered the study, reviewed the manuscript, and acquired financial support for the project leading this publication.

## SUPPLEMENTARY FIGURES

**Figure S1.**
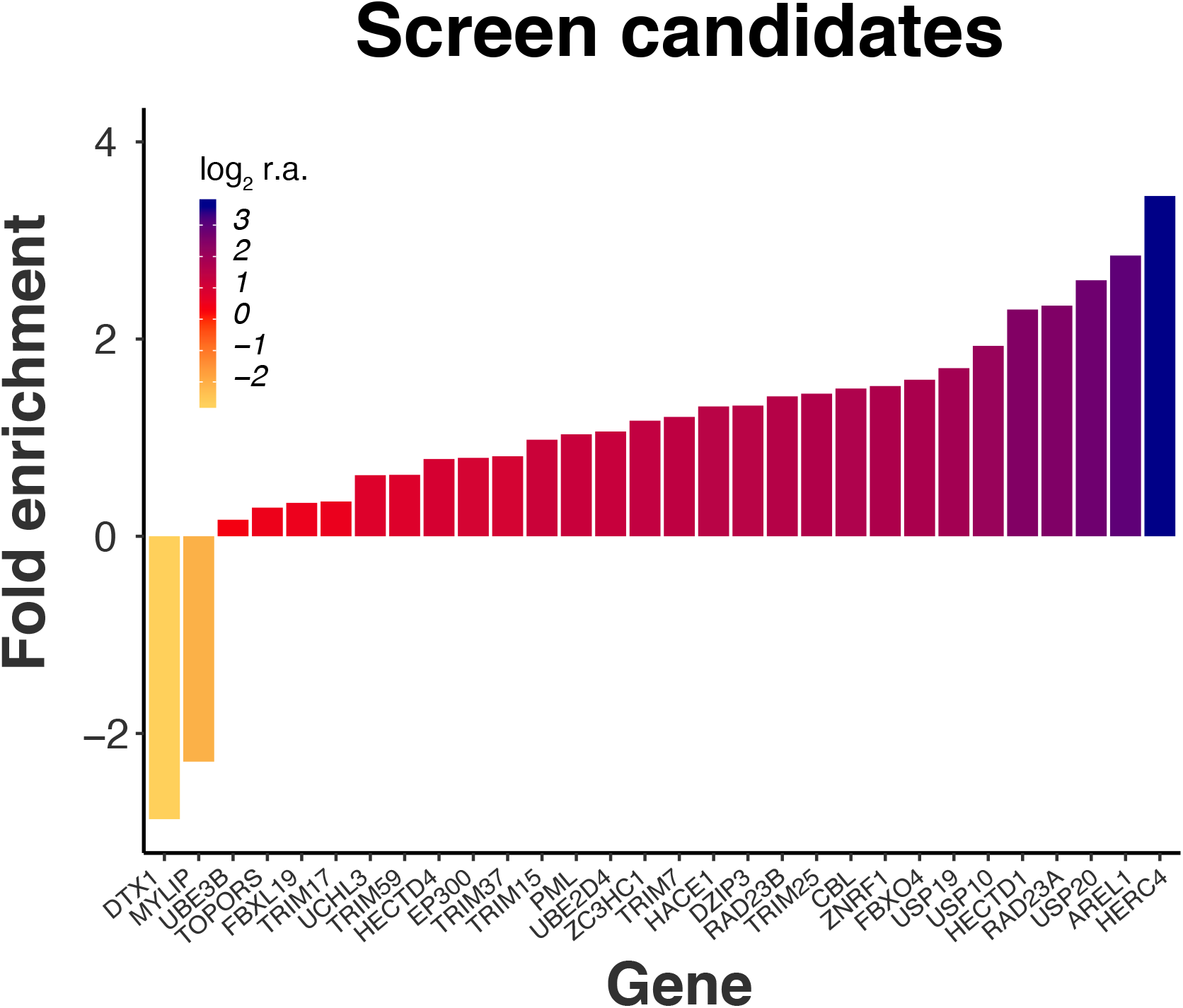
Candidate regulators of migration. We performed a loss-of-function screen using shRNAs targeting ubiquitination pathway-related genes in MDAMB231 cells. We implemented a cyclic functional selection and focused our analysis in cells that presented impaired migration. By sequencing DNA of selected cells, we calculated the frequency of each shRNA relative to the whole population of shRNAs after every cycle of selection. We then calculated the fold enrichment of each shRNA as their relative abundance in the fourth cycle compared to their abundance in the non-selected population of cells. The graph shows the mean fold enrichment (log_2_ of relative abundance [r.a.]) of at least two shRNAs targeting each candidate gene. Values above zero represent putative positive regulators of migration, and values below zero belong to shRNAs that were lost in the screen and could represent negative regulators of migration.

**Figure S2.**
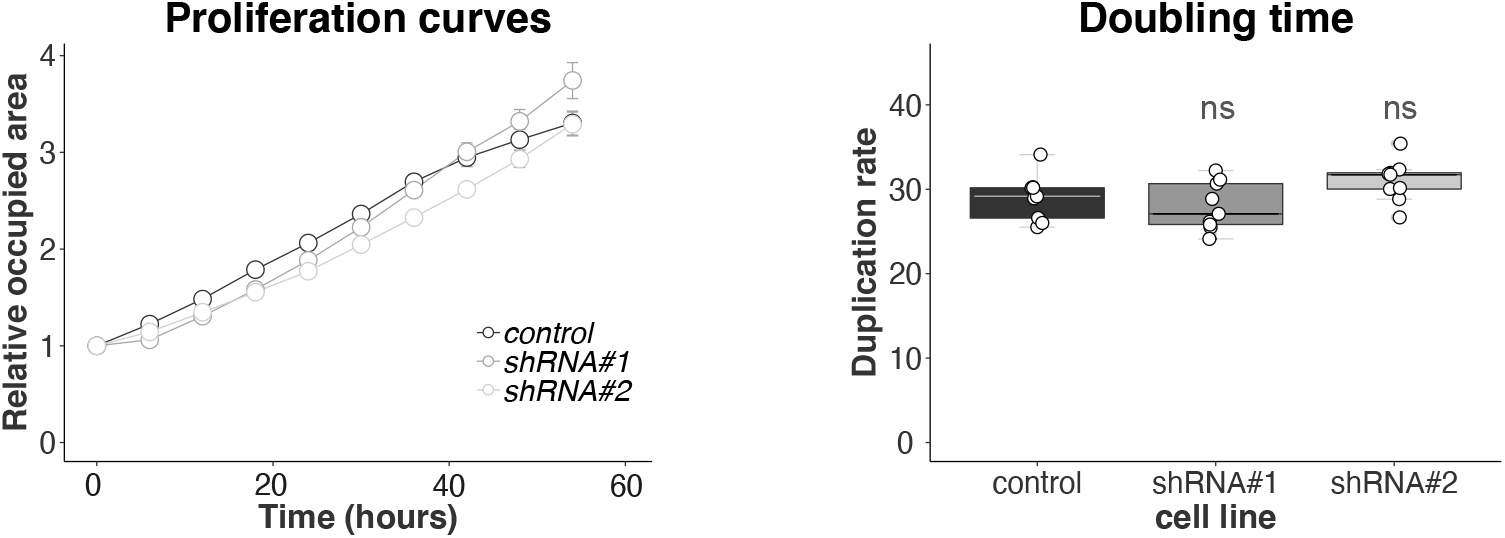
Doubling time of USP19 silenced MDAMB231 cells. An area-based analysis of proliferation rate was used to determine cell growth over time. Cells were seeded onto wells, allowed to attach and imaged at the indicated time points. Left: relative mean occupied area at each time point ± S.E., and right: doubling time calculated for control and USP19 silenced cell lines (n= 9, one-way ANOVA, Dunnett’s multiple comparison test. shRNA#1 p= 0.6284 and shRNA#2 p= 0.2123).

**Figure S3.**
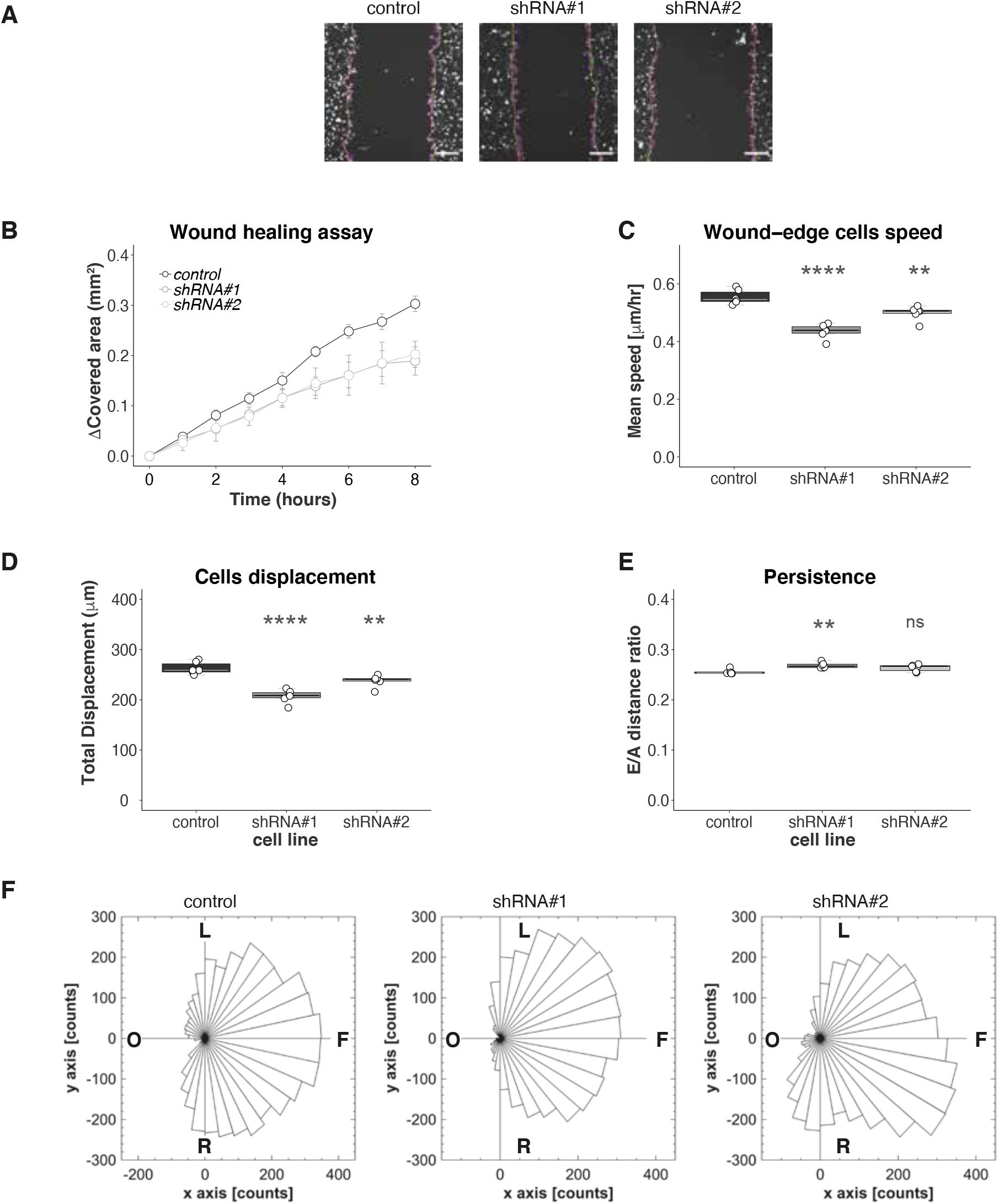
Detailed analysis on the wound healing assay in control or USP19 silenced MDAMB231 cells. We generated MDAMB231 cells expressing GFP tagged-H2B and then stably transduced them with control vector or shRNAs # 1 and # 2 directed against USP19. For the wound healing assays, a scratch was made with a pipette tip on a monolayer of the different cell cultures, and time-lapse imaging monitored the number of migrating cells across the border. (**A**) Representative images of a wound healing assay at time= 0 hours. Cells circled in magenta were considered for the analysis. Data corresponding to circled cells in the gap area were manually excluded from the analysis. Data was collected from three independent experiments for each cell line. Scale bar= 100 μm. (**B**) The graph shows the gap covered area (mm^2^) at the indicated time. (**C**) Cells mean speed from each wound-edge was calculated throughout the length of the experiment. (n=6, one-way ANOVA, Dunnett’s multiple comparison test. shRNA#1 p< 0.0001 and shRNA#2 p= 0.0028). (**D**) Mean accumulated track displacement (total displacement) for cells in each wound-edge was calculated until the end of the experiment (n=6, one-way ANOVA, Dunnett’s multiple comparison test. shRNA#1 p< 0.0001 and shRNA#2 p= 0.0059). (**E**) Persistence was calculated as the Euclidean/Accumulated track displacement ratio for control or USP19 silenced MDAMB231 cells at each wound-edge. (n=6, one-way ANOVA, Dunnett’s multiple comparison test. shRNA#1 p= 0.0003 and shRNA#2 p= 0.0527). (**F**) Rose diagrams representing the orientation of migration trajectories relative to cells position at time= 0 hours. Cells from the left wound-edge of three videos per cell line are shown, and each sector indicates the frequency of trajectory orientation. Control cell line (n= 131), shRNA#1 (n= 107) and shRNA#2 (n= 123). F: forward to direction of migration, L: left to direction of migration, O: opposite to direction of migration, R: right to direction of migration.

**Figure S4.**
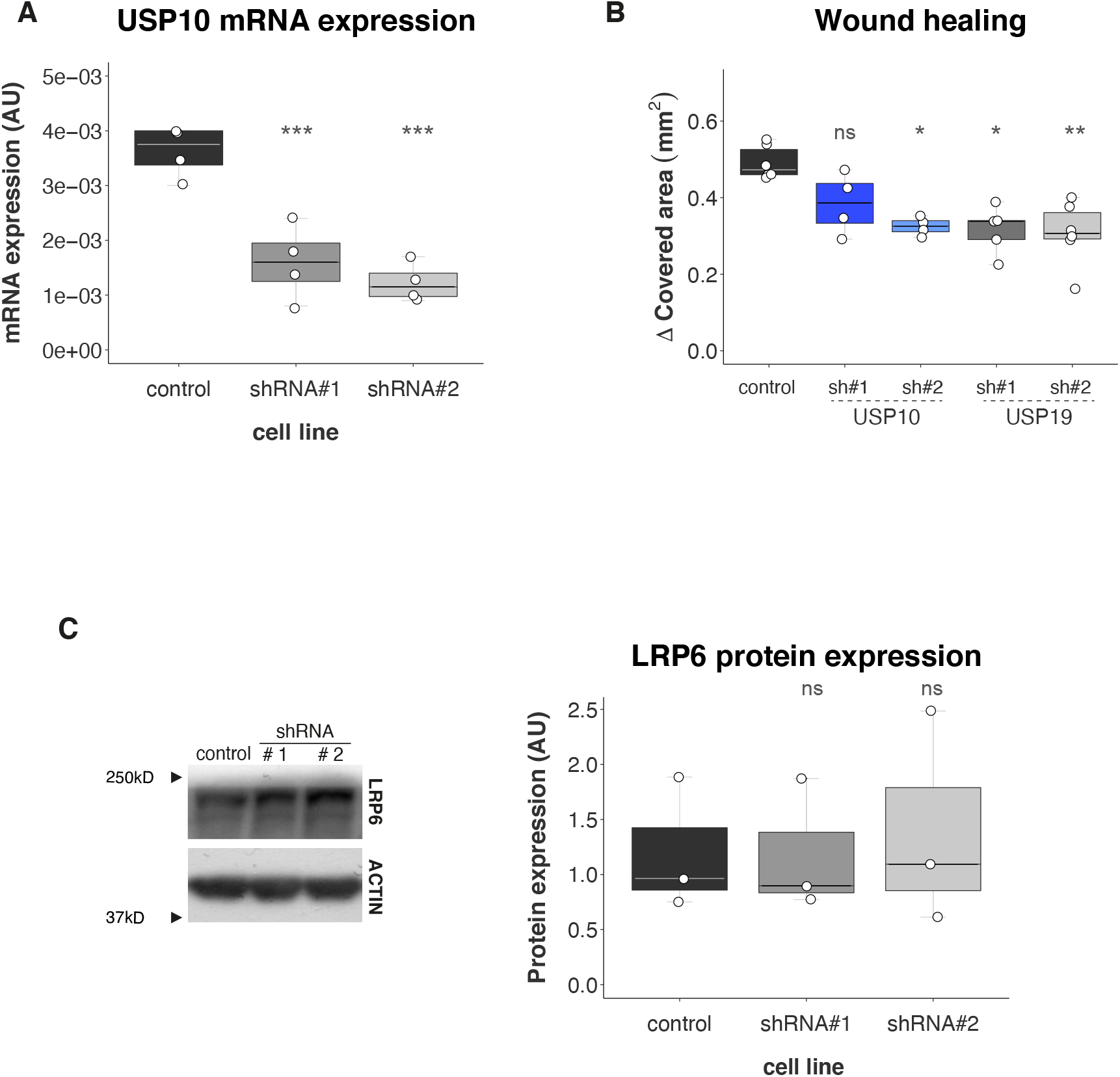
USP10 silencing effects in MDAMB231 cells migratory potential. We generated USP10 silenced MDAMB231 cells and compared the effect on migration with USP19 depletion experiments. (**A**) Efficiencies of shRNA-mediated USP10 knockdown were confirmed by RT-PCR (n= 4, one-way ANOVA, Dunnett’s multiple comparison test. shRNA#1 p= 0.0004 and shRNA#2 p= 0.0002, relative to mRNA expression in control cells). (**B**) Wound healing assays were used to analyze migration. The graph shows the gap covered area for control or cells stably expression shRNAs targeting USP10 or USP19 after 8 hours (n≥4 Kruskal-Wallis, Dunn’s multiple comparison test. USP10 shRNA#1 p= 0.4576, USP10 shRNA#2 p= 0.0397, USP19 shRNA#1 p= 0.0141 and USP19 shRNA#2 p= 0.0060). (**C**) LRP6 protein expression was evaluated in control or USP10 silenced MDAMB231 cells. Left: representative image of a blot; right: LRP6 signal quantification (n=3, Kruskal-Wallis and Dunn’s multiple comparison test. shRNA#1 p> 0.9999 and shRNA#2 p> 0.9999).

**Figure S5.**
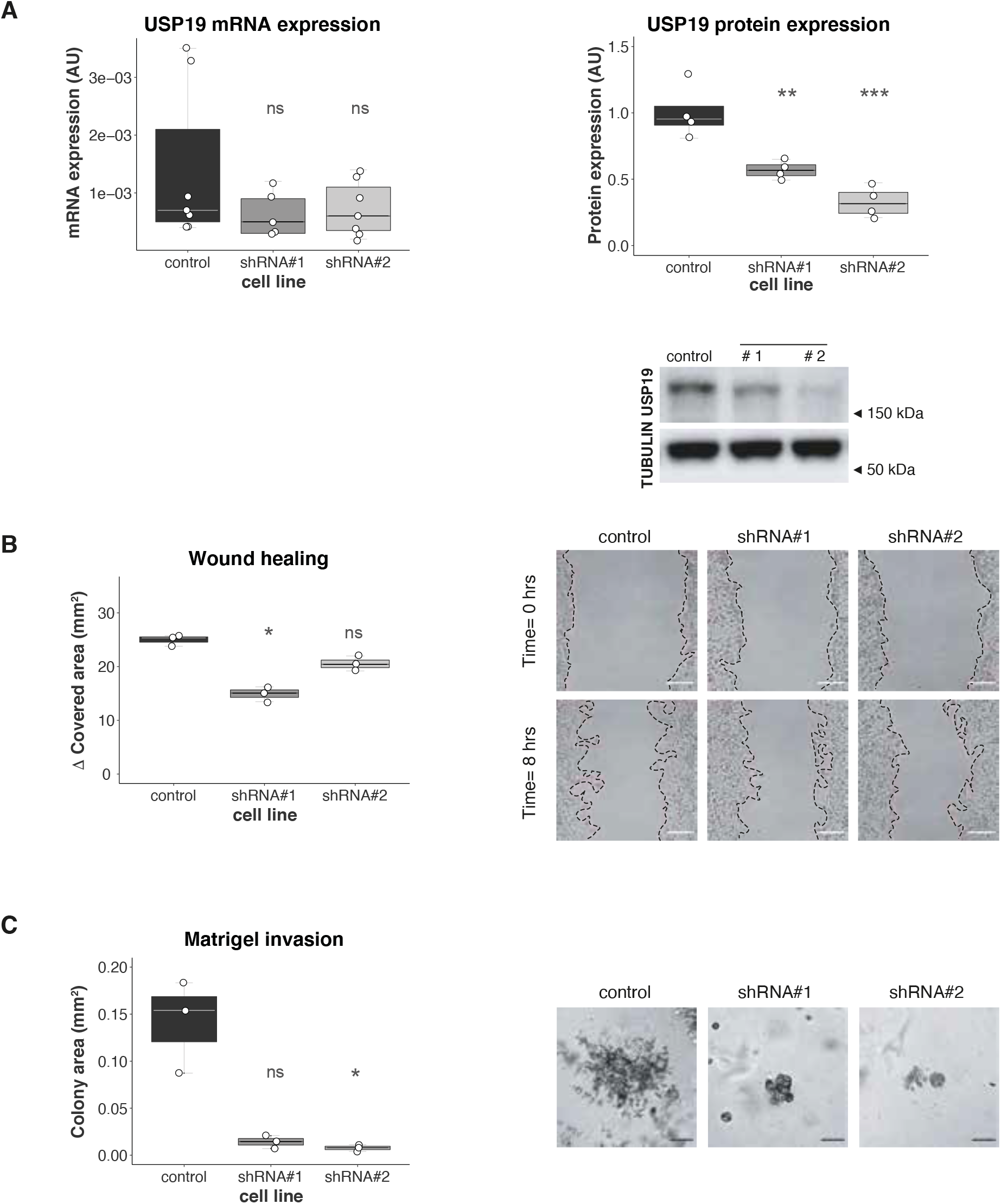
In vitro validation in MDAMB436 cells. USP19 silencing effects were studied in MDAMB436 cells. (**A**) Left: Efficiencies of shRNA-mediated target gene knockdown were confirmed by RT-PCR (n≥ 5, Kruskal-Wallis, Dunn’s multiple comparison test. shRNA#1 p= 0.6028 and shRNA#2 p= 0.8335) and Right: Western Blotting (n= 4, one-way ANOVA, Dunnett’s multiple comparison test. shRNA#1 p= 0.0039 and shRNA#2 p= 0.0002). (**B**) Cells migratory potential was evaluated by wound healing assay. Left: gap covered area (mm^2^) after 8 hours (n= 3, Kruskal-Wallis, Dunn’s multiple comparison test. shRNA#1 p= 0.0146 and shRNA#2 p= 0.3594); right: representative areas in a wound healing experiment at the indicated time points. Scale bar= 100 μm. (**C**) Invasiveness was assessed with Matrigel 3D experiments. Left: Colony area was calculated at the end of the experiment (n= 3, Kruskal-Wallis, Dunn’s multiple comparison test. shRNA#1 p= 0.2021 and shRNA#2 p= 0.0341); right: a representative area for the cell invasion assay after 5 days in culture is shown. Scale bar= 200 μm.

**Figure S6.**
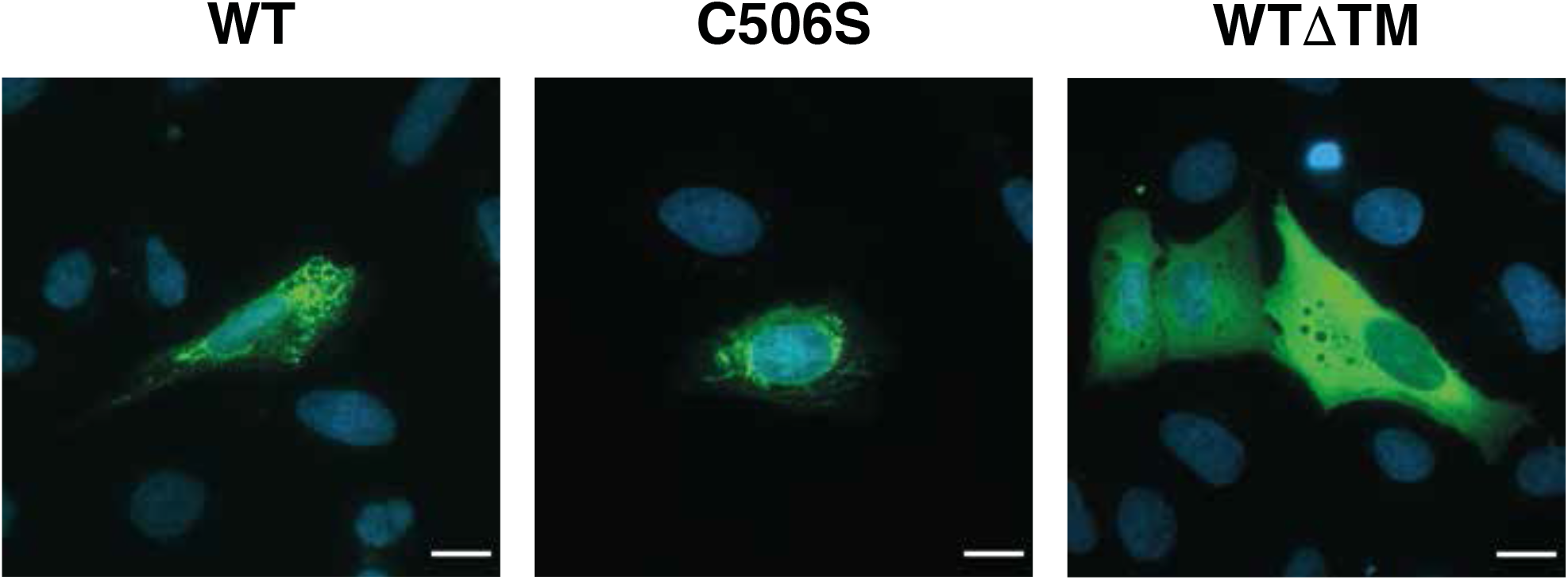
USP19 overexpression in U2OS cells. There are a variety of USP19 mRNA transcripts generated by alternative splicing, and not all of them show the same functional properties. In this regard, changes in the last exon coding sequence generate soluble isoforms and endoplasmic reticulum (ER) membrane-bound isoforms. In order to analyze whether USP19’s transmembrane domain was required for migration and 3D growth, we generated a GFP-tagged USP19 construct (wild type for its catalytic activity) with a point mutation that generates a premature stop codon. This mutation prevented the transmembrane domain to be translated, therefore generating a protein that mimicked USP19’s soluble isoform (named WTΔTM). We analyzed overexpressed-USP19 proteins subcellular localization by transiently transfecting U2OS cells with wild type (WT), catalytically dead mutant (C506S) or cytoplasmic (WTΔTM) GFP-tagged USP19 constructs. Cells were fixed with 4% paraformaldehyde, stained with DAPI and mounted. Images were captured using an inverted fluorescence microscopy. Scale bar= 100 μm.

**Figure S7.**
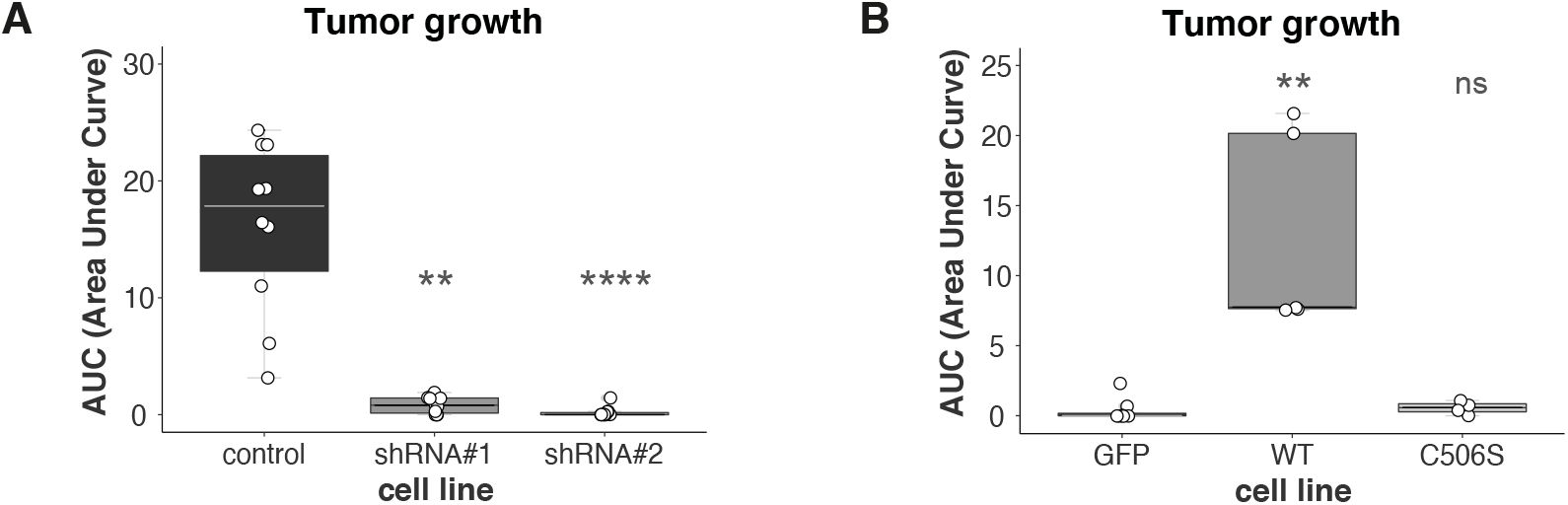
Tumor growth analysis. Tumor growth was analyzed by calculating the area under the tumor volume curves of mice injected with the different cell lines, at the end of the experiment. (**A**) AUC values of mice injected with MDAMB231 cells (n≥ 10, Kruskal-Wallis, Dunn’s multiple comparison test. shRNA#1 p= 0.0021 and shRNA#2 p< 0.0001). (**B**) AUC values of mice injected with MCF7 cells (n≥ 4, Kruskal-Wallis, Dunn’s multiple comparison test. WT p= 0.0015 and C506S p= 0.7582).

**Figure S8.**
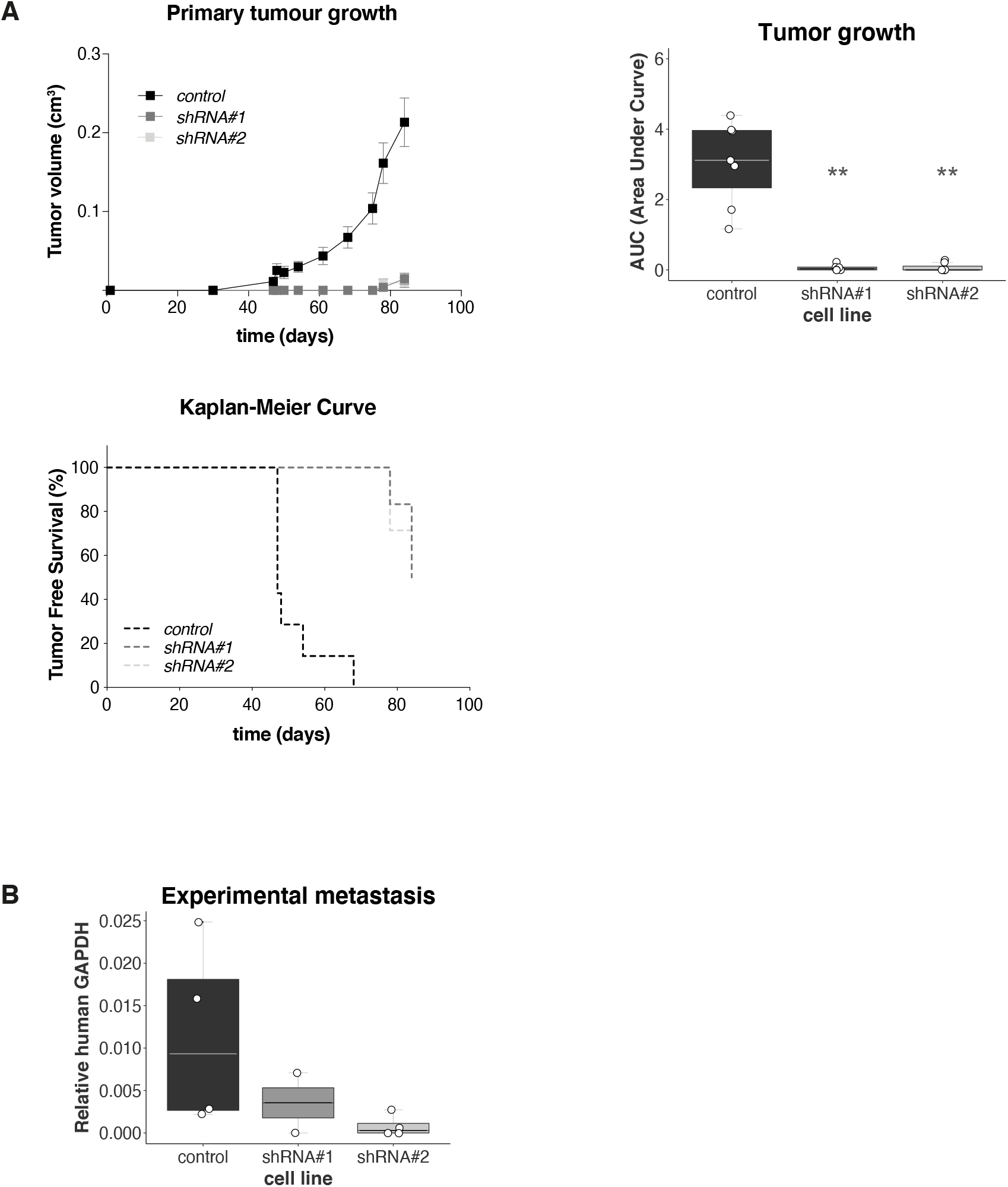
In vivo experiments with MDAMB436 cells. (**A**) Downregulation of USP19 attenuates tumorigenicity in vivo: control or USP19-silenced MDAMB436 cells were subcutaneously inoculated into the mammary fat pads of female NOD/SCID mice. Top left: Tumor volume was measured at the indicated time points (results show mean value ± S.E.). Top right: the area under the tumor volume curves was calculated at the end of the experiment (n≥ 6, Kruskal-Wallis, Dunn’s multiple comparison test. shRNA#1 p= 0.0053 and shRNA#2 p= 0.0014). Bottom: Kaplan-Meier curves were built for Tumor Free Survival (TFS) over time (n≥ 6, Log-Rank (Mantel-Cox) test, shRNA#1 p= 0.0003 and shRNA#2 p= 0.0001). (**B**) Experimental metastasis assay: NOD/SCID male mice were inoculated with MDAMB436 USP19-silenced cells through tail vein injection and after 2 months, lungs were harvested. Metastasis foci were estimated by qPCR human DNA quantification (n≥ 2).

**Figure S9.**
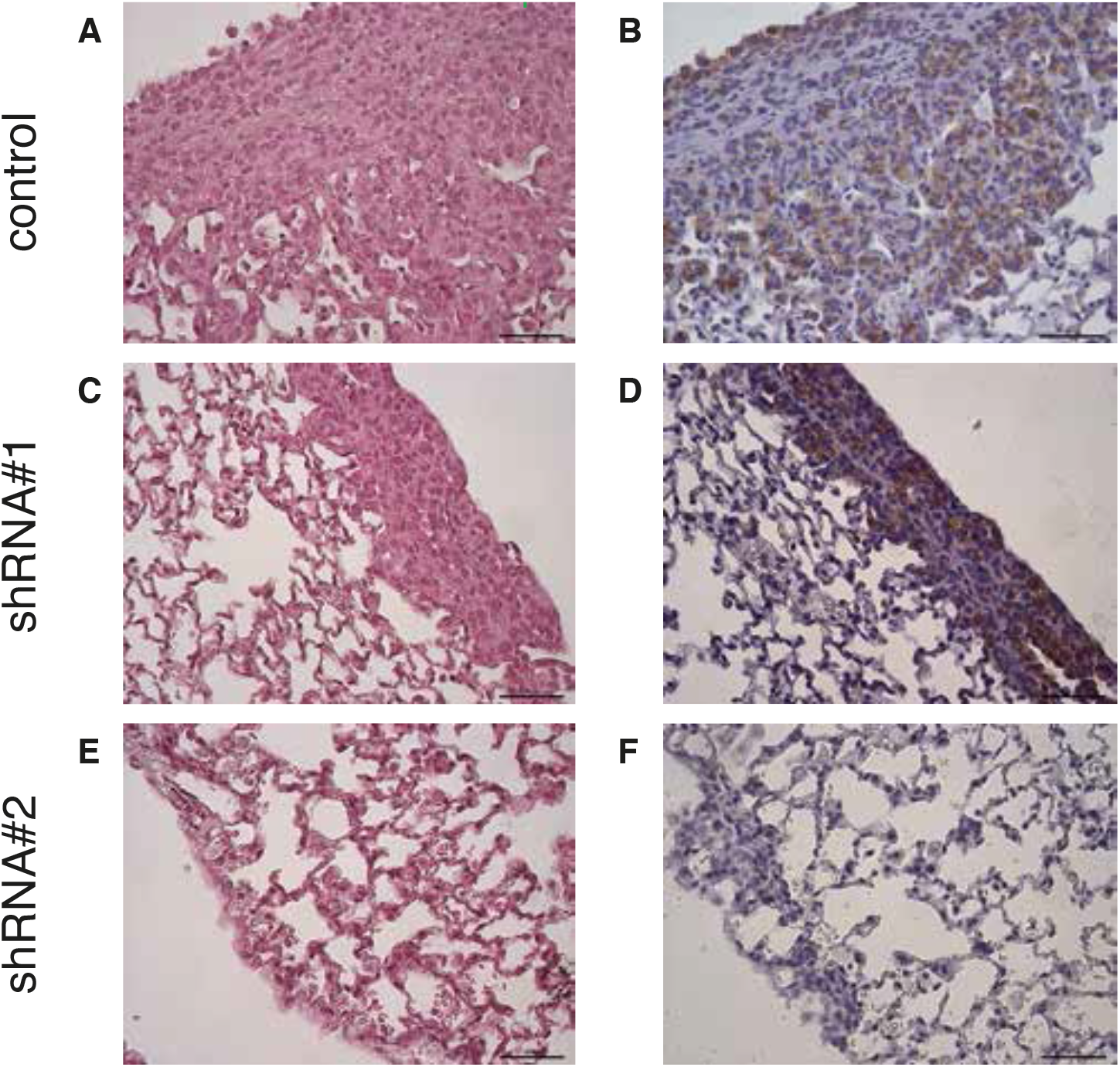
Metastatic load determination. Representative images of lung sections of mice injected with control or USP19-silenced MDAMB231 cells. Tissues were analyzed for metastatic load by Hematoxylin & Eosin staining (**A**, **C**, and **E**), and nodules tumor origin was confirmed by staining with the mouse anti-human CD44s monoclonal antibody (**B**, **D**, and **F**). Scale bar: 50 μm.

**Figure S10.**
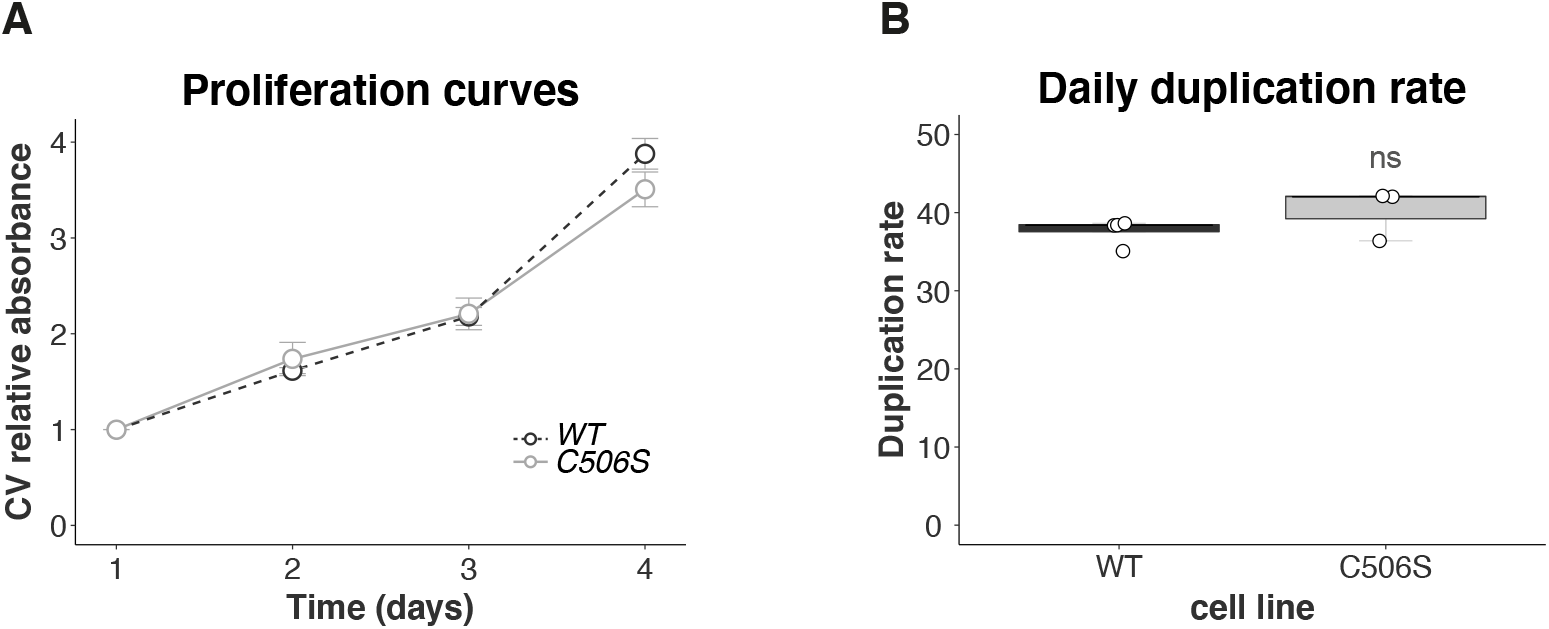
USP19 overexpression does not affect 2D proliferation in MCF7 cells. Crystal violet (CV) staining was used to determine cell growth over time. Cells were seeded onto wells and allowed to attach. At the indicated time points, cells were fixed and then stained at the end of the experiment. (**A**) The graph shows the mean relative CV absorbance every 24 hours. (**B**) Doubling time was calculated for MCF7 cells overexpressing WT or mutant C506S USP19 constructs (n≥ 3, Mann-Whitney test, p= 0.4000).

**Figure S11.**
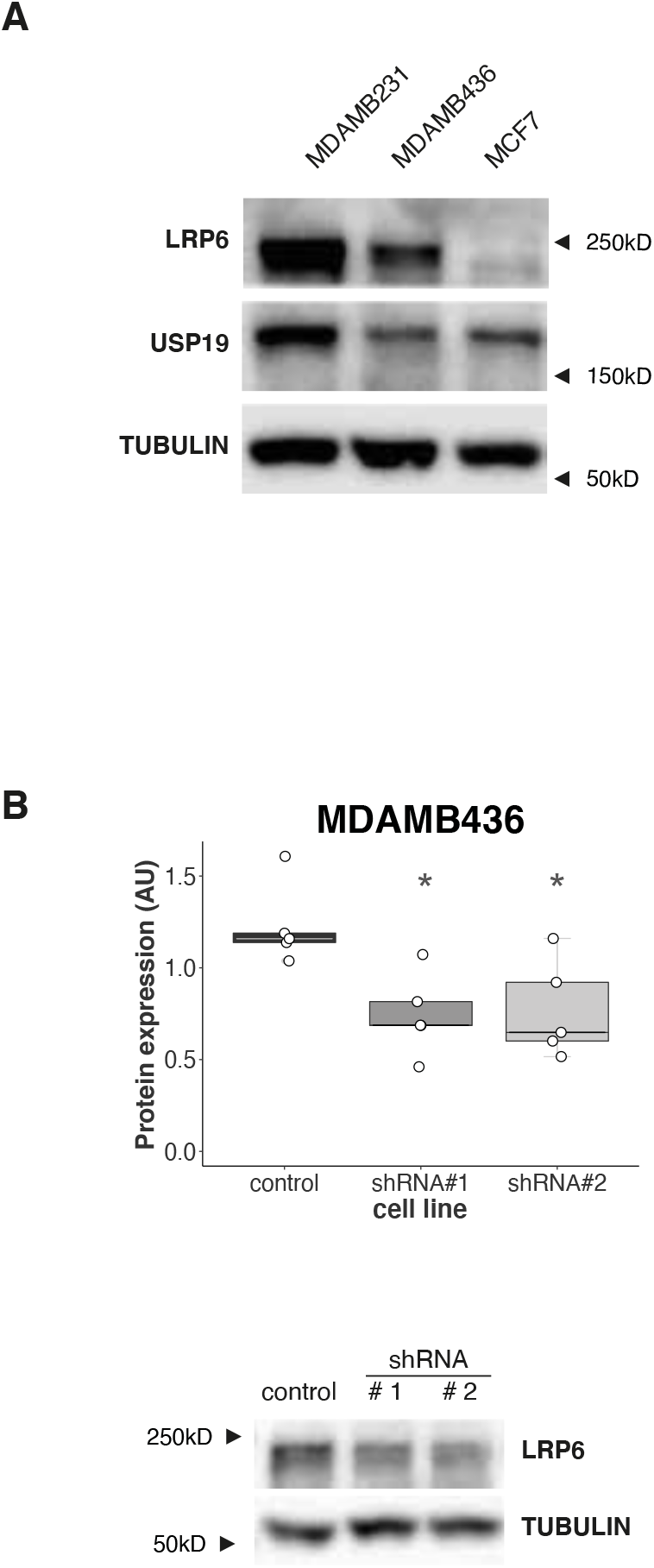
USP19 expression regulates LRP6 protein levels in breast cancer cell lines. (**A**) Western blotting was performed in order to analyze LRP6 protein expression in the breast cancer cell lines used in this manuscript. (**B**) LRP6 protein expression was evaluated in control or USP19 silenced MDAMB436 cells. Top: LRP6 signal quantification (n=5, one-way ANOVA, Dunnett’s multiple comparison test. shRNA#1 p= 0.0139 and shRNA#2 p= 0.0188), bottom: representative image of a blot.

**Figure S12.**
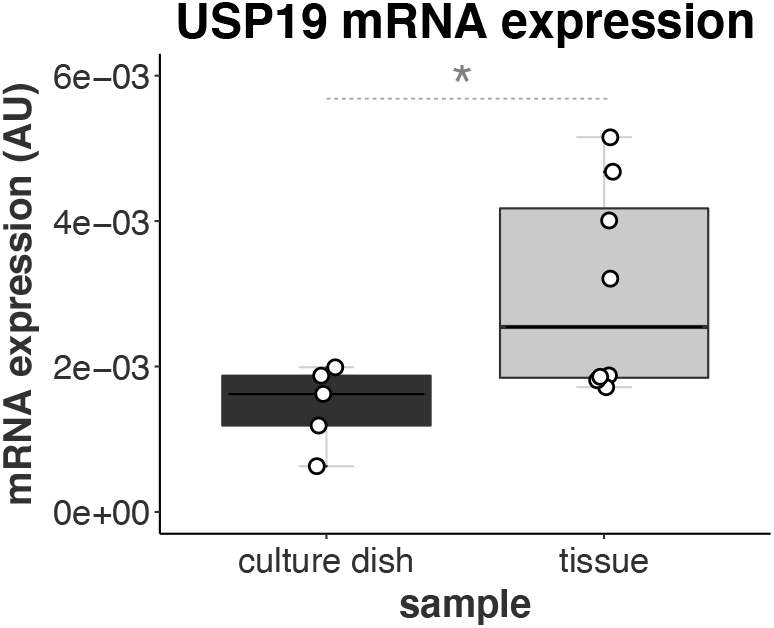
USP19 mRNA expression is increased in tissues. USP19 mRNA expression levels were analyzed by RT-qPCR in cultured MDAMB231 cells expressing PLKO.1 empty vector (growing in culture dishes), or in tumors that generated in NOD/SCID mice upon injection of MDAMB231 cells expressing PLKO.1 empty vector (n≥ 5, Mann-Whitney test, p= 0.0326).

**Figure S13.**
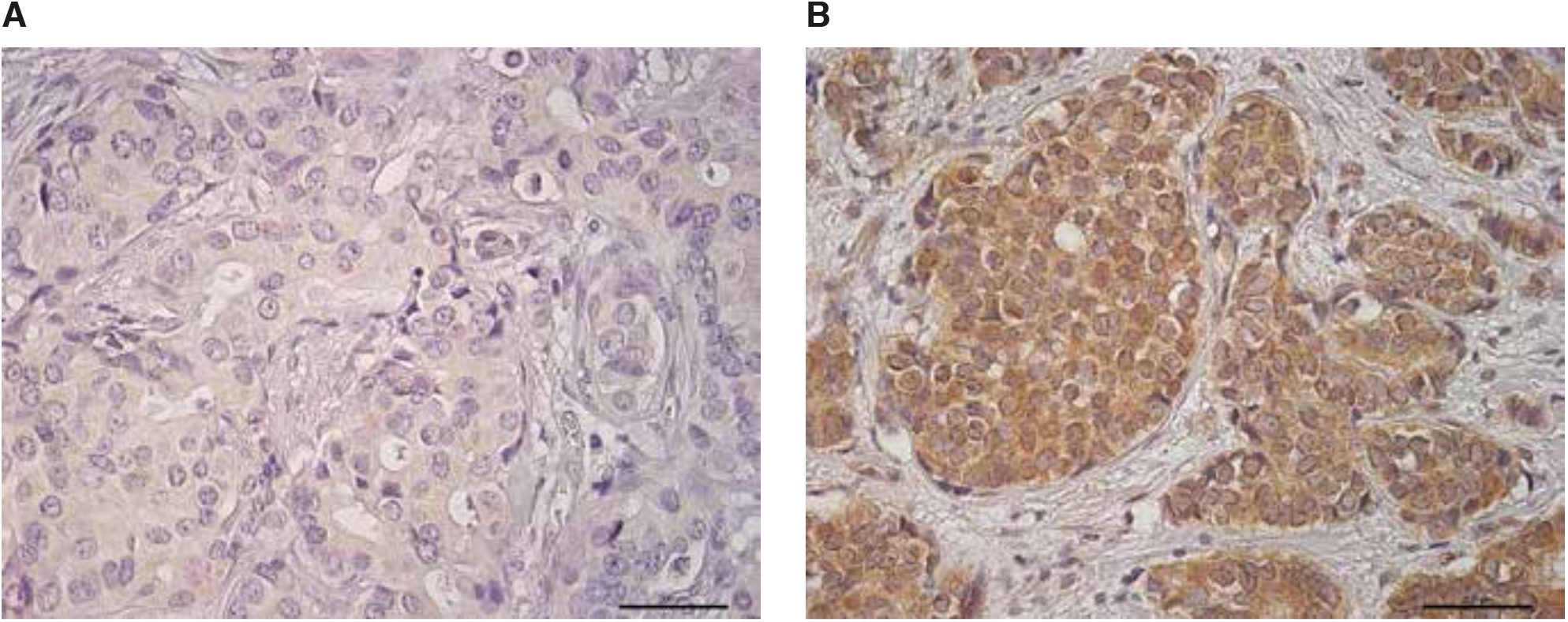
USP19 staining on FFPE breast cancer tissues. Immunohistochemical staining: examples of (**A**) low and (**B**) high expression of USP19 in breast cancer. Scale bar: 200 μm.

## SUPPLEMENTARY TABLES

**Table S1.** References for known regulators of migration in the candidate gene list. List of the candidate genes obtained after our screen, that had already been described in the literature as regulators of migration and the corresponding supporting references. The table does not review all the known functions of each already-published gene.

| GENE | REFERENCE Title | DOI |
| --- | --- | --- |
| DTX1 | Integrative computational analysis of transcriptional and epigenetic alterations implicates *DTX1* as a putative tumor suppressor gene in HNSCC. | 10.18632/oncotarget.14856 |
| MYLIP | miR-19b promotes breast cancer metastasis through targeting MYLIP and its related cell adhesion molecules. | 10.18632/oncotarget.19278 |
| FBXL19 | SCF E3 Ligase F-box Protein Complex SCF(FBXL19) Regulates Cell Migration by Mediating Rac1 Ubiquitination and Degradation. | 10.1096/fj.12-223099 |
|  | A new mechanism of RhoA ubiquitination and degradation: Roles of SCF<sup>FBXL19</sup> E3 ligase and Erk2. | 10.1016/j.bbamcr.2013.07.005 |
|  | F-box protein complex FBXL19 regulates TGFβ1-induced E-cadherin down-regulation by mediating Rac3 ubiquitination and degradation. | 10.1186/1476-4598-13-76 |
| UHL3 | UHL3 Promotes Pancreatic Cancer Progression and Chemo-Resistance Through FOXM1 Stabilization. | www.ajcr.us/ISSN:2156-6976/ajcr0089233 |
|  | UHL3 promotes ovarian cancer progression by stabilizing TRAF2 to activate the NF-κB pathway. | 10.1038/s41388-019-0987-z |
| TRIM59 | TRIM59 promotes breast cancer motility by suppressing p62-selective autophagic degradation of PDCD10. | 10.1371/journal.pbio.3000051. |
|  | TRIM59 loss in M2 macrophages promotes melanoma migration and invasion by upregulating MMP-9 and Madcam1. | 10.18632/aging.102351 |
|  | TRIM59 overexpression correlates with poor prognosis and contributes to breast cancer progression through AKT signaling pathway. | 10.1002/mc.22897 |
|  | An TRIM59-CDK6 axis regulates growth and metastasis of lung cancer. | 10.1111/jcmm.14052 |
|  | TRIM59 Promotes the Proliferation and Migration of Non-Small Cell Lung Cancer Cells by Upregulating Cell Cycle Related Proteins. | 10.1371/journal.pone.0142596 |
| HECTD4 | LncRNA HEIH promotes cell proliferation, migration and invasion in cholangiocarcinoma by modulating miR-98-5p/HECTD4. | 10.1016/j.biopha.2020.109916. |
| EP300 | EP300 as an oncogene correlates with poor prognosis in esophageal squamous carcinoma. | 10.7150/jca.34261. |
|  | The p300/YY1/miR-500a-5p/HDAC2 signalling axis regulates cell proliferation in human colorectal cancer. | 10.1038/s41467-018-08225-3. |
|  | P300 promotes proliferation, migration, and invasion via inducing epithelial-mesenchymal transition in non-small cell lung cancer cells. | 10.1186/s12885-018-4559-3. |
| PML | Ubiquitination of tumor suppressor PML regulates prometastatic and immunosuppressive tumor microenvironment. | 10.1172/JCI89957. |
|  | PML represses lung cancer metastasis by suppressing the nuclear EGFR-mediated transcriptional activation of MMP2. | 10.4161/15384101.2014.94921. |
| TRIM37 | TRIM37 promoted the growth and migration of the pancreatic cancer cells. | 10.1007/s13277-015-4078-7. |
|  | Knockdown of TRIM37 suppresses the proliferation, migration and invasion of glioma cells through the inactivation of PI3K/Akt signaling pathway. | 10.1016/j.biopha.2018.01.054. |
|  | TRIM37 promotes cell invasion and metastasis by regulating SIP1-mediated epithelial-mesenchymal transition in gastric cancer. | 10.2147/OTT.S178446. |
|  | Over-expression of TRIM37 promotes cell migration and metastasis in hepatocellular carcinoma by activating Wnt/β-catenin signaling. | 10.1016/j.bbrc.2015.07.089. |
|  | TRIM37 promotes epithelial-mesenchymal transition in colorectal cancer. | 10.3892/mmr.2017.6125. |
| TRIM15 | High Expression of TRIM15 Is Associated with Tumor Invasion and Predicts Poor Prognosis in Patients with Gastric Cancer. | 10.1080/08941939.2019.1705443. |
| HACE1 | Loss of HACE1 promotes colorectal cancer cell migration via upregulation of YAP1. | 10.1002/jcp.27653. |
|  | Overexpression of HACE1 in gastric cancer inhibits tumor aggressiveness by impeding cell proliferation and migration. | 10.1002/cam4.1496. |
|  | The tumour suppressor HACE1 controls cell migration by regulating Rac1 degradation. | 10.1038/onc.2012.189. |
| CBL | The E3 ligase C-CBL inhibits cancer cell migration by neddylation of the proto-oncogene c-Src. | 10.1038/s41388-018-0354-5. |
|  | c-CBL regulates melanoma proliferation, migration, invasion and the FAK-SRC-GRB2 nexus. | 10.18632/oncotarget.10861. |
|  | c-Cbl regulates migration of v-Abl-transformed NIH 3T3 fibroblasts via Rac1. | 10.1016/j.yexcr.2005.03.010. |
| | c-Cbl regulates αPix-mediated cell migration and invasion. | 10.1016/j.bbrc.2014.10.129. |
| TRIM25 | TRIM25 blockade by RNA interference inhibited migration and invasion of gastric cancer cells through TGF-β signaling. | 10.1038/srep19070. |
|  | The ubiquitin ligase TRIM25 inhibits hepatocellular carcinoma progression by targeting metastasis associated 1 protein. | 10.1002/iub.1661. |
|  | Overexpression of TRIM25 in Lung Cancer Regulates Tumor Cell Progression. | 10.1177/1533034615595903. |
| | Tripartite motif containing 25 promotes proliferation and invasion of colorectal cancer cells through TGF-β signaling. | 10.1042/BSR20170805. |
| USP19 | Ubiquitin Specific Peptidase 19 Is a Prognostic Biomarker and Affect the Proliferation and Migration of Clear Cell Renal Cell Carcinoma | 10.3892/or.2020.7565 |
| USP10 | USP10 promotes proliferation and migration and inhibits apoptosis of endometrial stromal cells in endometriosis through activating the Raf-1/MEK/ERK pathway. | 10.1152/ajpcell.00272.2018. |
|  | USP10 regulates the stability of the EMT-transcription factor Slug/SNAI2. | 10.1016/j.bbrc.2018.05.156. |
|  | Deubiquitinating enzyme USP10 promotes hepatocellular carcinoma metastasis through deubiquitinating and stabilizing Smad4 protein. | 10.1002/1878-0261.12596. |
| HECTD1 | The E3 Ubiquitin Ligase HectD1 Suppresses EMT and Metastasis by Targeting the +TIP ACF7 for Degradation. | 10.1016/j.celrep.2017.12.096. |
|  | HECTD1 regulates the expression of SNAIL: Implications for epithelial-mesenchymal transition. | 10.3892/ijo.2020.5002. |
| RAD23A (HR23A) | HR23A-knockdown lung cancer cells exhibit epithelial-to-mesenchymal transition and gain stemness properties through increased Twist1 stability. | 10.1016/j.bbamcr.2019.118537. |
| USP20 | USP20 positively regulates tumorigenesis and chemoresistance through β-catenin stabilization. | 10.1038/s41418-018-0138-z. |
| HERC4 | HERC4 Is Overexpressed in Hepatocellular Carcinoma and Contributes to the Proliferation and Migration of Hepatocellular Carcinoma Cells. | 10.1089/dna.2016.3626. |
|  | RNA interference of HERC4 inhibits proliferation, apoptosis and migration of cervical cancer Hela cells | 10.3969/j.issn.1673-4254.2017.02.15. |
|  | A miRNA-HERC4 pathway promotes breast tumorigenesis by inactivating tumor suppressor LATS1. | 10.1007/s13238-019-0607-2. |

**Table S2.**
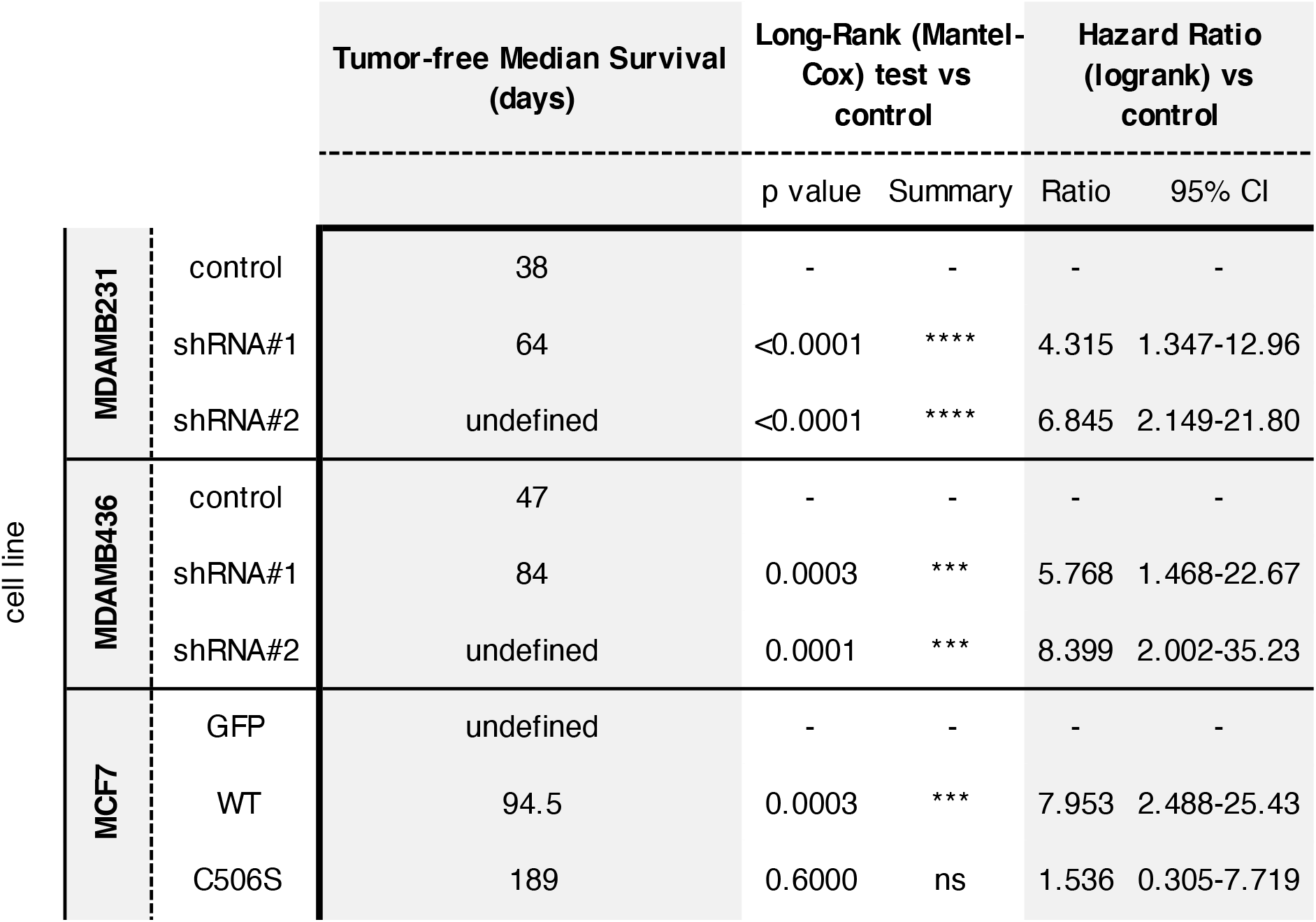
Analysis of Tumor Free Survival curves in USP19 silencing and overexpression in vivo experiments. USP19 effects on tumor growth were analyzed by injecting genetically manipulated cell lines in the mammary fat pad of NOD/SCID mice and measuring tumor growth over time. For each control or USP19 silenced cell line, in the case of MDAMB231 or MDAMB436 cells, and for each control or USP19 overexpressing cell line, in the case of MCF7 cells, tumor growth curves were built. Using this data, we calculated median tumor-free survival time (expressed in days) and performed Long-Rank (Mantel-Cox) tests of each cell line relative to their corresponding control. Hazard ratio ranks relative to control treatments are also indicated in the table.

**Table S3.** E-cadherin and USP19 staining in human biopsies. Detailed description of E-cadherin and USP19 staining in the patients’ breast cancer biopsies used in this study.

| N. | BIOPSY N# | CODE | AGE | FOLLOW-UP<br>(months) | E-caderin | E-cad_int | USP19_C% | USP19_C_int |
| --- | --- | --- | --- | --- | --- | --- | --- | --- |
| 213 | 83/7127 | RM 1 | 58.7 | 298 | 98 | 3 | 0 | 0 |
| 214 | 92/720 | RM 2 | 40.7 | 165 | 84 | 3 | 82 | 2 |
| 215 | 96/1827 | RM 3 | 61.7 | 149 | 80 | 3 | 84 | 2 |
| 219 | 92/2237 | RM 7 | 45.6 | 188 | 75 | 3 | 71 | 2 |
| 220 | 96/1511 | RM 8 | 59.7 | 114 | 97 | 3 | 85 | 2 |
| 221 | 96/70 | RM 9 | 75.8 | 146 | 16 | 2 | 11 | 2 |
| 223 | 95/2705 | RM 11 | 59.5 | 148 | 100 | 3 | 0 | 0 |
| 225 | 96/561 | RM 13 | 56.2 | 149 | 0 | 0 | 0 | 0 |
| 226 | 95/4980 | RM 14 | 38.6 | 108 | 51 | 2 | 0 | 0 |
| 227 | 95/3549 | RM 15 | 56.2 | 156 |  |  | 88 | 2 |
| 228 | 93/1074 | RM 16 | 60.2 | 170 | 96 | 3 | 0 | 0 |
| 232 | 95/3029 | RM 20 | 44.1 | 158 | 18 | 2 | 100 | 3 |
| 234 | 94/499 | RM 22 | 55.3 | 175 | 42 | 2 | 98 | 2 |
| 235 | 96/350 | RM 23 | 61.1 | 99 | 87 | 3 | 68 | 2 |
| 240 | 96/4647 | RM 28 | 64.1 | 100 |  |  | 0 | 0 |
| 241 | 96/5756 | RM 29 | 45.6 | 132 | 81 | 3 | 92 | 2 |
| 242 | 94/3629 | RM 30 | 70.5 | 153 | 35 | 2 | 6 | 2 |
| 244 | 96/3170 | RM 32 | 73.5 | 99 | 41 | 2 | 0 | 0 |
| 246 | 96/5503 | RM 34 | 49.8 | 143 | 54 | 2 | 43 | 2 |
| 247 | 94/3573 | RM 35 | 65.2 | 134 | 5 | 1 | 66 | 2 |
| 251 | 95/2532 | RM 39 | 48.7 | 155 | 71 | 2 | 94 | 2 |
| 254 | 96/4664,<br>4662/6/7 | RM 42 | 37.3 | 131 | 37 | 2 | 0 | 0 |
| 256 | 95/1794X | RM 44 | 45.5 | 160 |  |  | 60 | 2 |
| 258 | 96/2810 | RM 46 | 66.7 | 89 | 71 | 2 | 72 | 2 |
| 260 | 96/5222 | RM 48 | 43.3 | 110 | 62 | 2 | 95 | 3 |
| 261 | 94/3682 | RM 49 | 37.6 | 111 | 0 | 0 | 0 | 0 |
| 262 | 94/1319 | RM 50 | 55.5 | 171 | 77 | 3 | 88 | 2 |
| 264 | 97/4510 | RM 52 | 50.2 | 129 | 13 | 2 | 0 | 0 |
| 265 | 97/3300 | RM 53 | 58.6 | 77 | 87 | 3 | 71 | 2 |
| 267 | 97/1147 | RM 55 | 55.7 | 135 | 4 | 1 | 69 | 2 |
| 268 | 97/5339 | RM 56 | 41.2 | 132 | 16 | 2 | 9 | 1 |
| 274 | 97/3554 | RM 62 | 72.8 | 95 | 5 | 1 | 100 | 2 |
| 280 | 97/2971 | RM 68 | 67.3 | 130 | 82 | 3 | 0 | 0 |
| 281 | 97/3389 | RM 69 | 51.5 | 135 | 3 | 1 | 80 | 2 |
| 285 | 97/2638 | RM 73 | 61.6 | 133 | 2 | 1 | 0 | 0 |
| 288 | 98/3968 | RM 76 | 42.1 | 117 | 4 | 1 | 0 | 0 |
| 290 | 98/4874 | RM 78 | 40.8 | 122 | 9 | 1 | 9 | 2 |
| 291 | 98/2370 | RM 79 | 39.6 | 75 | 0 | 0 | 66 | 2 |
| 294 | 98/7174 | RM 82 | 63.6 | 118 | 85 | 3 | 0 | 0 |
| 295 | 98/3768 | RM 83 | 63.0 | 106 | 47 | 2 | 0 | 0 |
| 298 | 98/9072 | RM 86 | 64.6 | 117 | 100 | 3 | 0 | 0 |
| 300 | 98/3737 | RM 88 | 66.0 | 88 | 77 | 3 | 0 | 0 |
| 301 | 98/1642 | RM 89 | 67.0 | 77 | 79 | 3 | 8 | 2 |
| 303 | 98/8533 | RM 91 | 44.3 | 101 | 6 | 1 | 0 | 0 |
| 305 | 98/3165 | RM 93 | 66.0 | 112 | 0 | 0 | 0 | 0 |
| 306 | 98/1566 | RM 94 | 61.6 | 123 | 4 | 1 | 4 | 2 |
| 311 | 98/1187 | RM 99 | 41.3 | 125 | 21 | 2 | 63 | 2 |
| 313 | 99/11766 | RM 101 | 53.2 | 107 | 8 | 1 | 0 | 0 |
| 314 | 99/10805 | RM 102 | 51.9 | 104 | 97 | 3 | 0 | 0 |
| 316 | 99/2464 | RM 104 | 48.0 | 37 | 5 | 1 | 0 | 0 |
| 317 | 99/793 | RM 105 | 46.9 | 113 | 0 | 0 | 41 | 2 |
| 319 | 99/11878 | RM 106 | 56.3 | 57 | 0 | 0 | 62 | 2 |
| 320 | 99/10246 | RM 107 | 50.6 | 68 | 74 | 3 | 0 | 0 |
| 322 | 99/1994 | RM 109 | 51.9 | 86 | 46 | 2 | 40 | 2 |
| 323 | 99/795 | RM 110 | 46.9 | 82 | 0 | 0 | 68 | 3 |
| 325 | 99/10154 | RM 112 | 42.1 | 55 | 89 | 3 | 0 | 0 |
| 326 | 99/4615 | RM 113 | 44.9 | 107 | 33 | 2 | 0 | 0 |
| 327 | 99/1579 | RM 114 | 70.6 | 79 | 26 | 2 | 0 | 0 |
| 328 | 99/794 | RM 115 | 46.9 | 82 | 3 | 1 | 7 | 2 |
| 330 | 99/10091 | RM 117 | 33.9 | 108 | 0 | 0 | 0 | 0 |
| 331 | 99/4087 | RM 118 | 56.7 | 77 | 41 | 2 | 0 | 0 |
| 332 | 99/1378 | RM 119 | 37.3 | 115 | 48 | 2 | 0 | 0 |
| 333 | 99/707 | RM 120 | 59.4 | 49 | 6 | 1 | 61 | 2 |
| 334 | 99/11246 | RM 121 | 51.8 | 104 | 2 | 1 | 0 | 0 |
| 335 | 99/9569 | RM 122 | 65.1 | 107 | 3 | 1 | 0 | 0 |
| 336 | 99/2441 | RM 123 | 68.3 | 85 | 95 | 3 | 0 | 0 |
| 339 | 00/2106 | RM 126 | 35.6 | 98 | 9 | 1 | 4 | 2 |
| 341 | 00/1325/2 | RM 128 | 66.3 | 60 |  |  | 64 | 1 |
| 343 | 99/13089 | RM 130 | 56.7 | 68 | 0 | 0 | 69 | 1 |
| 345 | 00/3002/1 | RM 132 | 62.1 | 60 |  |  | 0 | 0 |
| 346 | 00/1182 | RM 133 | 65.6 | 101 | 0 | 0 | 98 | 3 |
| 347 | 99/13376 | RM 134 | 66.8 | 104 | 81 | 3 | 36 | 1 |
| 348 | 99/12969 | RM 135 | 55.6 | 99 | 85 | 3 | 0 | 0 |
| 349 | 00/4188 | RM 136 | 72.3 | 33 | 77 | 3 | 0 | 0 |
| 350 | 00/2579 | RM 137 | 62.6 | 99 | 8 | 1 | 100 | 3 |
| 351 | 00/467 | RM 138 | 61.1 | 103 | 92 | 3 | 100 | 3 |
| 352 | 99/13331 | RM 139 | 53.6 | 66 | 0 | 0 | 0 | 0 |
| 353 | 99/12422/2 | RM 140 | 39.0 | 82 |  |  | 0 | 0 |
| 354 | 00/4225 | RM 141 | 67.1 | 97 | 70 | 3 | 0 | 0 |
| 355 | 00/1798 | RM 142 | 66.1 | 43 | 97 | 3 | 0 | 0 |
| 356 | 00/395 | RM 143 | 61.2 | 99 | 6 | 1 | 94 | 2 |
| 358 | 99/12089 | RM 145 | 59.6 | 101 | 90 | 3 | 0 | 0 |
| 360 | 00/1521/5 | RM 147 | 43.6 | 103 |  |  | 0 | 0 |
| 361 | 00/26 | RM 148 | 73.4 | 97 | 29 | 2 | 0 | 0 |
| 362 | 99/12324/1 | RM 149 | 68.0 | 69 |  |  | 0 | 0 |
| 364 | 00/468 | RM 151 | 53.9 | 70 | 94 | 3 | 0 | 0 |
| 365 | 00/3295 | RM 152 | 76.0 | 45 | 96 | 3 | 0 | 0 |
| 366 | 00/4998 | RM 153 | 38.9 | 94 | 6 | 1 | 0 | 0 |
| 367 | 00/5256 | RM 154 | 46.1 | 91 | 80 | 3 | 0 | 0 |
| 368 | 00/7139 | RM 155 | 61.5 | 91 | 2 | 1 | 46 | 1 |
| 369 | 00/1570 | RM 156 | 53.8 | 52 | 100 | 3 | 0 | 0 |
| 370 | 00/5073 | RM 157 | 46.6 | 49 | 8 | 1 | 0 | 0 |
| 371 | 00/4999 | RM 158 | 61.6 | 46 | 2 | 1 | 72 | 2 |
| 372 | 00/7167 | RM 159 | 37.8 | 94 | 3 | 1 | 95 | 2 |
| 373 | 00/5082 | RM 160 | 64.9 | 99 | 5 | 1 | 0 | 0 |
| 375 | 00/3394 | RM 162 | 67.2 | 90 | 9 | 1 | 100 | 2 |
| 376 | 00/5119 | RM 163 | 40.9 | 98 | 12 | 2 | 90 | 2 |
| 377 | 00/7215 | RM 164 | 69.6 | 94 | 11 | 2 | 100 | 3 |
| 378 | 00/1610 | RM 165 | 66.5 | 98 |  |  | 0 | 0 |
| 379 | 00/1957/7 | RM 166 | 75.0 | 48 |  |  | 87 | 2 |
| 380 | 00/3625 | RM 167 | 56.8 | 53 | 10 | 2 | 69 | 2 |
| 381 | 00/5192 | RM 168 | 43.9 | 90 | 7 | 1 | 91 | 2 |
| 382 | 00/7240 | RM 169 | 59.3 | 89 | 0 | 0 | 72 | 2 |
| 385 | 00/2617 | RM 172 | 74.6 | 56 | 81 | 3 | 13 | 1 |
| 386 | 00/4397 | RM 173 | 45.4 | 100 |  |  | 94 | 2 |
| 387 | 00/5234 | RM 174 | 32.8 | 54 | 5 | 1 | 91 | 2 |
| 389 | 99/2946 | RM 176 | 55.2 | 107 |  |  | 0 | 0 |
| 391 | 01/2658 | RM 178 | 51.5 | 34 |  |  | 52 | 1 |
| 392 | 01/7932 | RM 179 | 58.4 | 82 | 0 | 0 | 85 | 1 |
| 394 | 01/6229 | RM 181 | 40.4 | 6 | 0 | 0 | 0 | 0 |
| 397 | 01/7712 | RM 184 | 50.3 | 77 | 81 | 3 | 0 | 0 |
| 398 | 01/7037 | RM 185 | 57.7 | 82 | 26 | 2 | 0 | 0 |
| 399 | 01/5438 | RM 186 | 82.9 | 39 | 4 | 1 | 0 | 0 |
| 400 | 01/3265 | RM 187 | 44.6 | 83 | 59 | 2 | 0 | 0 |
| 403 | 01/6840 | RM 190 | 52.9 | 81 | 2 | 1 | 0 | 0 |
| 404 | 01/5320 | RM 191 | 68.2 | 29 | 10 | 2 | 0 | 0 |
| 407 | 01/7617/2 | RM 194 | 47.1 | 79 |  |  | 0 | 0 |
| 408 | 01/6580 | RM 195 | 56.7 | 82 | 0 | 0 | 7 | 1 |
| 409 | 01/4931/1 | RM 196 | 59.1 | 86 |  |  | 8 | 2 |
| 411 | 01/2037/7 | RM 198 | 66.6 | 63 |  |  | 61 | 2 |
| 412 | 01/7581 | RM 199 | 68.4 | 41 | 86 | 3 | 0 | 0 |
| 413 | 01/6307/1 | RM 200 | 55.2 | 42 |  |  | 0 | 0 |
| 414 | 01/3414/1 | RM 201 | 67.1 | 88 |  |  | 0 | 0 |
| 416 | 01/693 | RM 203 | 55.6 | 89 | 90 | 3 | 0 | 0 |
| 418 | 01/2037 | RM 205 | 66.4 | 57 |  |  | 0 | 0 |
| 419 | 01/4291 | RM 206 | 73.4 | 83 | 0 | 0 | 0 | 0 |
| 420 | 01/7 | RM 207 | 54.3 | 81 | 93 | 3 | 17 | 2 |
| 421 | 01/887 | RM 208 | 44.3 | 89 | 3 | 1 | 100 | 2 |
| 425 | 01/160 | RM 212 | 31.2 | 78 | 9 | 1 | 4 | 1 |
| 426 | 01/946 | RM 213 | 64.6 | 51 | 0 | 0 | 93 | 2 |
| 428 | 01/3458 | RM 215 | 55.8 | 83 | 77 | 3 | 0 | 0 |
| 429 | 01/4128 | RM 216 | 35.7 | 82 | 16 | 2 | 0 | 0 |
| 430 | 01/155 | RM 217 | 64.1 | 93 | 6 | 1 | 5 | 1 |
| 431 | 01/1268 | RM 218 | 51.4 | 92 | 46 | 2 | 0 | 0 |
| 433 | 01/5091 | RM 220 | 72.1 | 82 | 100 | 3 | 4 | 1 |
| 435 | 01/3192 | RM 222 | 50.9 | 53 | 4 | 1 | 5 | 1 |
| 436 | 01/1275/2 | RM 223 | 75.0 | 73 |  |  | 0 | 0 |
| 437 | 01/1936 | RM 224 | 39.0 | 77 | 6 | 1 | 90 | 2 |
| 439 | 01/3739 | RM 226 | 76.0 | 75 | 5 | 1 | 54 | 2 |
| 442 | 98/1603 | RM 229 | 59.6 | 46 | 86 | 3 | 0 | 0 |
| 443 | 96/404 | RM 230 | 50.7 | 12 | 18 | 2 | 0 | 0 |
| 444 | 92/1605 | RM 231 | 53.3 | 94 |  |  | 0 | 0 |
| 446 | 93/3375 | RM 233 | 65.1 | 90 | 8 | 1 | 0 | 0 |
| 447 | 97/4228 | RM 234 | 82.9 | 57 | 2 | 1 | 100 | 2 |
| 450 | 96/4480 | RM 237 | 50.6 | 12 | 85 | 3 | 9 | 1 |
| 451 | 90/1158 | RM 238 | 66.6 | 158 | 5 | 1 | 57 | 2 |
| 455 | 91/2410 | RM 242 | 45.5 | 144 |  |  | 98 | 2 |
| 457 | 96/3988 | RM 244 | 47.0 | 67 | 100 | 3 | 97 | 3 |
| 458 | 95/5063 | RM 245 | 72.7 | 23 | 76 | 3 | 71 | 2 |
| 459 | 91/1312 | RM 246 | 39.7 | 93 | 5 | 1 | 96 | 2 |
| 460 | 94/2465 | RM 247 | 61.5 | 135 | 28 | 2 | 61 | 1 |
| 461 | 00/3494 | RM 248 | 76.6 | 11 | 0 | 0 | 91 | 3 |
| 462 | 96/169 | RM 249 | 45.2 | 47 | 91 | 3 | 100 | 3 |
| 463 | 93/4502 | RM 250 | 54.4 | 42 | 35 | 2 | 83 | 2 |
| 464 | 87/8972 | RM 251 | 36.2 | 121 | 2 | 1 | 95 | 2 |
| 465 | 00/5709 | RM 252 | 68.4 | 73 | 0 | 0 | 0 | 0 |
| 466 | 97/6196 | RM 253 | 70.4 | 98 | 94 | 3 | 0 | 0 |
| 467 | 94/2414 | RM 254 | 45.6 | 166 | 0 | 0 | 16 | 1 |
| 469 | 00/2106/4 | RM 256 | 35.3 | 166 |  |  | 47 | 2 |
| 470 | 97/4512 | RM 257 | 58.1 | 78 |  |  | 0 | 0 |
| 471 | 90/2010 | RM 258 | 65.9 | 166 |  |  | 4 | 2 |
| 473 | 99/2622 | RM 260 | 42.7 | 43 | 95 | 3 | 0 | 0 |
| 475 | 98/1991 | RM 262 | 63.6 | 120 | 47 | 2 | 0 | 0 |
| 477 | 91/2728 | RM 264 | 61.9 | 110 | 7 | 1 | 0 | 0 |
| 479 | 96/4304 | RM 266 | 73.8 | 94 | 84 | 3 | 93 | 2 |
| 482 | 98/5936 | RM 269 | 55.9 | 14 | 8 | 1 | 93 | 2 |
| 483 | 98/6391 | RM 270 | 67.8 | 24 | 10 | 2 | 55 | 2 |
| 485 | 98/328 | RM 272 | 76.0 | 40 |  |  | 0 | 0 |

**Table S4.** Characterization of patient cohort used in this manuscript. 168 N0 patients with T1/T2 tumors with primary unilateral breast carcinoma from the Regina Elena National Cancer Institute, Rome, Italy were included in this study. Patients age at diagnosis, menopausal status and outcome are indicated in the table. Moreover, their tumors characteristics are stated, including molecular subtype, tumor size, histological type, grade, estrogen receptor (ER) status, progesterone receptor (PR) status, Ki-67 status, Human Epidermal Growth Factor Receptor 2 (HER2) status and USP19 staining level, as absolute and percentual values.

| Variable | Value (%) |
| --- | --- |
| **Age at diagnosis (yr)** |  |
| Median | 56.7 |
| <50 | 51 (30.4) |
| 50-65 | 69 (41.1) |
| >65 | 48 (28.6) |
| **Menopausal status** |  |
| Pre/perimenopausal | 54 (32.1) |
| Postmenopausal | 114 (67.9) |
| **Molecular subtypes** |  |
| Luminal A | 71 (42.3) |
| Luminal B | 57 (33.9) |
| HER2 positive | 8 (4.8) |
| Triple negative | 32 (19.0) |
| **Tumor size** |  |
| ≤ 2 cm | 109 (64.9) |
| > 2 cm | 59 (35.1) |
| **Histotypes** |  |
| Ductal carcinoma | 135 (80.4) |
| Lobular carcinoma | 24 (14.3) |
| Other | 9 (5.3) |
| **Tumor grade** |  |
| 1 | 21 (12.5) |
| 2-3 | 147 (87.5) |
| **ER** |  |
| Negative | 43 (25.6) |
| Positive | 125 (74.4) |
| **PR** |  |
| Negative | 59 (35.1) |
| Positive | 109 (64.9) |
| **Ki-67** |  |
| Negative | 90 (53.6) |
| Positive | 78 (46.4) |
| **HER2** |  |
| Negative | 149 (88.7) |
| Positive | 19 (11.3) |
| **USP19** |  |
| Low | 106 (63.1) |
| High | 62 (36.9) |
| **Patient outcome** |  |
| Without recurrence | 119 (70.8) |
| Local recurrence | 20 (11.9) |
| Distant recurrence | 29 (17.3) |

**Table S5.** Characterization of USP19 protein expression in human samples. The table shows USP19 status according to clinicopathological features of patients, including tumor size and grade, estrogen receptor (ER), progesterone receptor (PR) and Human Epidermal Growth Factor Receptor 2 (HER2) status, subtype and ki-67 staining (Pearson’s χ^2^ test). Cases are indicated as absolute and percentual values.

| Variable | USP19 |  | P* |
| --- | --- | --- | --- |
|  | Low: n (%) | High: n (%) |  |
| Tumor size |  |  |  |
| ≤ 2 cm | 74 (69.8) | 35 (56.5) | 0.095 |
| > 2 cm | 32 (30.2) | 27 (42.5) |  |
| Tumor grade |  |  |  |
| 1 | 17 (16.0) | 4 (6.5) | 0.091 |
| 2-3 | 89 (84.0) | 58 (93.5) |  |
| ER |  |  |  |
| Negative | 27 (25.5) | 16 (25.8) | 1,000 |
| Positive | 79 (74.5) | 46 (74.2) |  |
| PR |  |  |  |
| Negative | 40 (37.7) | 19 (30.6) | 0.404 |
| Positive | 66 (62.3) | 43 (69.4) |  |
| Ki-67 |  |  |  |
| Negative | 61 (57.5) | 29 (46.8) | 0.201 |
| Positive | 45 (42.5) | 33 (53.2) |  |
| HER2 |  |  |  |
| Negative | 100 (94.3) | 49 (79.0) | 0.003* |
| Positive | 6 (5.7) | 13 (21.0) |  |
| Subtype |  |  |  |
| Luminal A | 48 (67.6) | 23 (32.4) | 0.014* |
| Luminal B | 31 (54.4) | 26 (45.6) |  |
| HER2 | 2 (25.0) | 6 (75.0) |  |
| TN | 25 (63.1) | 7 (36.9) |  |

## REFERENCES

[1] Trepat X, Chen Z, Jacobson K. Cell migration. Compr Physiol 2012;2:2369–92.

[2] Vicente-Manzanares M, Horwitz AR. Cell migration: an overview. Methods Mol Biol 2011;769:1–24.

[3] Acloque H, Adams MS, Fishwick K, Bronner-Fraser M, Nieto MA. Epithelial-mesenchymal transitions: the importance of changing cell state in development and disease. The Journal of clinical investigation 2009;119:1438–49.

[4] Yang J, Weinberg RA. Epithelial-mesenchymal transition: at the crossroads of development and tumor metastasis. Dev Cell 2008;14:818–29.

[5] Hanahan D, Weinberg RA. Hallmarks of cancer: the next generation. Cell 2011;144:646–74.

[6] Gupta GP, Massague J. Cancer metastasis: building a framework. Cell 2006;127:679–95.

[7] Lefranc F, Brotchi J, Kiss R. Possible future issues in the treatment of glioblastomas: special emphasis on cell migration and the resistance of migrating glioblastoma cells to apoptosis. J Clin Oncol 2005;23:2411–22.

[8] Megalizzi V, Mathieu V, Mijatovic T, Gailly P, Debeir O, De Neve N, et al. 4-IBP, a sigma1 receptor agonist, decreases the migration of human cancer cells, including glioblastoma cells, *in vitro* and sensitizes them *in vitro* and *in vivo* to cytotoxic insults of proapoptotic and proautophagic drugs. Neoplasia 2007;9:358–69.

[9] Wells A, Grahovac J, Wheeler S, Ma B, Lauffenburger D. Targeting tumor cell motility as a strategy against invasion and metastasis. Trends Pharmacol Sci 2013;34:283–9.

[10] Palmer T, Ashby W, Lewis J, Zijlstra A. Targeting tumor cell motility to prevent metastasis. Adv Drug Deliv Rev 2012;63:568–81.

[11] Eckhardt BL, Francis PA, Parker BS, Anderson RL. Strategies for the discovery and development of therapies for metastatic breast cancer. Nature reviews Drug discovery 2012;11:479–97.

[12] Ciechanover A, Hod Y, Hershko A. A heat-stable polypeptide component of an ATP-dependent proteolytic system from reticulocytes. Biochem Biophys Res Commun 1978;81:1100–5.

[13] Hershko A, Ciechanover A, Heller H, Haas A, Rose IA. Proposed role of ATP in protein breakdown: Conjugation of proteins with multiple chains of the polypeptide of ATP-dependent proteolysis. Proc Natl Acad Sci U S A 1980;77:1783–6.

[14] Husnjak K, Dikic I. Ubiquitin-Binding Proteins: Decoders of Ubiquitin-Mediated Cellular Functions. Annual Review of Biochemistry 2012;81:291–322.

[15] Pickart CM. Mechanisms underlying ubiquitination. Annual Review of Biochemistry 2001;70:503–33.

[16] Nijman SMB, Luna-Vargas MPA, Velds A, Brummelkamp TR, Dirac AMG, Sixma TK, et al. A genomic and functional inventory of deubiquitinating enzymes. Cell 2005;123:773–86.

[17] Schaefer A, Nethe M, Hordijk PL. Ubiquitin links to cytoskeletal dynamics, cell adhesion and migration. The Biochemical journal 2012;442:13–25.

[18] Mohr S, Bakal C, Perrimon N. Genomic screening with RNAi: results and challenges. Annu Rev Biochem 2010;79:37–64.

[19] Ryan AASaTE. RNAi Screening for the Discovery of Novel Modulators of Human Disease. Current Pharmaceutical Biotechnology 2010:735–56.

[20] Mullenders J, Bernards R. Loss-of-function genetic screens as a tool to improve the diagnosis and treatment of cancer. Oncogene 2009;28:4409–20.

[21] So RWL, Chung SW, Lau HHC, Watts JJ, Gaudette E, Al-Azzawi ZAM, et al. Application of CRISPR genetic screens to investigate neurological diseases. Mol Neurodegener 2019;14:41.

[22] Gerhards NM, Rottenberg S. New tools for old drugs: Functional genetic screens to optimize current chemotherapy. Drug Resist Updat 2018;36:30–46.

[23] Yao H, He G, Yan S, Chen C, Song L, Rosol TJ, et al. Triple-negative breast cancer: is there a treatment on the horizon? Oncotarget 2017;8:1913–24.

[24] Garrido-Castro AC, Lin NU, Polyak K. Insights into Molecular Classifications of Triple-Negative Breast Cancer: Improving Patient Selection for Treatment. Cancer Discov 2019;9:176–98.

[25] Lee KL, Kuo YC, Ho YS, Huang YH. Triple-Negative Breast Cancer: Current Understanding and Future Therapeutic Breakthrough Targeting Cancer Stemness. Cancers (Basel) 2019;11.

[26] Mehanna J, Haddad FG, Eid R, Lambertini M, Kourie HR. Triple-negative breast cancer: current perspective on the evolving therapeutic landscape. Int J Womens Health 2019;11:431–7.

[27] He WT, Zheng XM, Zhang YH, Gao YG, Song AX, van der Goot FG, et al. Cytoplasmic Ubiquitin-Specific Protease 19 (USP19) Modulates Aggregation of Polyglutamine-Expanded Ataxin-3 and Huntingtin through the HSP90 Chaperone. PLoS One 2016;11:e0147515.

[28] Wiles B, Miao M, Coyne E, Larose L, Cybulsky AV, Wing SS. USP19 deubiquitinating enzyme inhibits muscle cell differentiation by suppressing unfolded-protein response signaling. Molecular biology of the cell 2015;26:913–23.

[29] Hassink GC, Zhao B, Sompallae R, Altun M, Gastaldello S, Zinin NV, et al. The ER-resident ubiquitin-specific protease 19 participates in the UPR and rescues ERAD substrates. EMBO Rep 2009;10:755–61.

[30] Lee JG, Kim W, Gygi S, Ye Y. Characterization of the deubiquitinating activity of USP19 and its role in endoplasmic reticulum-associated degradation. The Journal of biological chemistry 2014;289:3510–7.

[31] He WT, Xue W, Gao YG, Hong JY, Yue HW, Jiang LL, et al. HSP90 recognizes the N-terminus of huntingtin involved in regulation of huntingtin aggregation by USP19. Sci Rep 2017;7:14797.

[32] Perrody E, Abrami L, Feldman M, Kunz B, Urbe S, van der Goot FG. Ubiquitin-dependent folding of the Wnt signaling coreceptor LRP6. eLife 2016;5.

[33] Bilir B, Kucuk O, Moreno CS. Wnt signaling blockage inhibits cell proliferation and migration, and induces apoptosis in triple-negative breast cancer cells. Journal of translational medicine 2013;11:12.

[34] Geyer FC, Lacroix-Triki M, Savage K, Arnedos M, Lambros MB, MacKay A, et al. beta-Catenin pathway activation in breast cancer is associated with triple-negative phenotype but not with CTNNB1 mutation. Modern pathology : an official journal of the United States and Canadian Academy of Pathology, Inc 2011;24:209–31.

[35] Khramtsov AI, Khramtsova GF, Tretiakova M, Huo D, Olopade OI, Goss KH. Wnt/beta-catenin pathway activation is enriched in basal-like breast cancers and predicts poor outcome. The American journal of pathology 2010;176:2911–20.

[36] Matsuda Y, Schlange T, Oakeley EJ, Boulay A, Hynes NE. WNT signaling enhances breast cancer cell motility and blockade of the WNT pathway by sFRP1 suppresses MDA-MB-231 xenograft growth. Breast Cancer Res 2009;11:R32.

[37] Raisch J, Cote-Biron A, Rivard N. A Role for the WNT Co-Receptor LRP6 in Pathogenesis and Therapy of Epithelial Cancers. Cancers (Basel) 2019;11.

[38] Chen Q, Hang Y, Zhang T, Tan L, Li S, Jin Y. USP10 promotes proliferation and migration and inhibits apoptosis of endometrial stromal cells in endometriosis through activating the Raf-1/MEK/ERK pathway. Am J Physiol Cell Physiol 2018;315:C863–C72.

[39] Ouchida AT, Kacal M, Zheng A, Ambroise G, Zhang B, Norberg E, et al. USP10 regulates the stability of the EMT-transcription factor Slug/SNAI2. Biochem Biophys Res Commun 2018;502:429–34.

[40] Yuan T, Chen Z, Yan F, Qian M, Luo H, Ye S, et al. Deubiquitinating enzyme USP10 promotes hepatocellular carcinoma metastasis through deubiquitinating and stabilizing Smad4 protein. Mol Oncol 2020;14:197–210.

[41] Hanahan D, Weinberg RA. Biological hallmarks of cancer. 9th ed: Wiley Blackwell; 2017.

[42] Welch DR, Hurst DR. Defining the Hallmarks of Metastasis. Cancer Res 2019;79:3011–27.

[43] Combaret L, Adegoke OA, Bedard N, Baracos V, Attaix D, Wing SS. USP19 is a ubiquitin-specific protease regulated in rat skeletal muscle during catabolic states. Am J Physiol Endocrinol Metab 2005;288:E693–700.

[44] Jin S, Tian S, Chen Y, Zhang C, Xie W, Xia X, et al. USP19 modulates autophagy and antiviral immune responses by deubiquitinating Beclin-1. EMBO J 2016;35:866–80.

[45] Gierisch ME, Pedot G, Walser F, Lopez-Garcia LA, Jaaks P, Niggli FK, et al. USP19 deubiquitinates EWS-FLI1 to regulate Ewing sarcoma growth. Sci Rep 2019;9:951.

[46] Lu Y, Adegoke OA, Nepveu A, Nakayama KI, Bedard N, Cheng D, et al. USP19 deubiquitinating enzyme supports cell proliferation by stabilizing KPC1, a ubiquitin ligase for p27Kip1. Mol Cell Biol 2009;29:547–58.

[47] Lim K, Choi J, Park J, Cho H, Park J, Lee E, et al. Ubiquitin specific protease 19 involved in transcriptional repression of retinoic acid receptor by stabilizing CORO2A. Oncotarget 2016;7:34759–72.

[48] Yamaguchi H, Condeelis J. Regulation of the actin cytoskeleton in cancer cell migration and invasion. Biochimica et biophysica acta 2007;1773:642–52.

[49] Nair NU, Das A, Rogkoti VM, Fokkelman M, Marcotte R, de Jong CG, et al. Migration rather than proliferation transcriptomic signatures are strongly associated with breast cancer patient survival. Sci Rep 2019;9:10989.

[50] Weathington NM, Mallampalli RK. Emerging therapies targeting the ubiquitin proteasome system in cancer. The Journal of clinical investigation 2014;124:6–12.

[51] Veggiani G, Gerpe MCR, Sidhu SS, Zhang W. Emerging drug development technologies targeting ubiquitination for cancer therapeutics. Pharmacol Ther 2019;199:139–54.

[52] Morrow JK, Lin H-K, Sun S-C, Zhang S. Targeting ubiquitination for cancer therapies. Future Medicinal Chemistry 2015;7:2333–50.

[53] Huang X, Dixit VM. Drugging the undruggables: exploring the ubiquitin system for drug development. Cell Research 2016;26:484–98.

[54] Cui J, Jin S, Wang RF. The BECN1-USP19 axis plays a role in the crosstalk between autophagy and antiviral immune responses. Autophagy 2016;12:1210–1.

[55] Gu Z, Shi W, Zhang L, Hu Z, Xu C. USP19 suppresses cellular type I interferon signaling by targeting TRAF3 for deubiquitination. Future Medicine 2017.

[56] Lei CQ, Wu X, Zhong X, Jiang L, Zhong B, Shu HB. USP19 Inhibits TNF-alpha- and IL-1beta-Triggered NF-kappaB Activation by Deubiquitinating TAK1. J Immunol 2019;203:259–68.

[57] Lim K-H, Choi J-H, Park J-H, Cho H-J, Park J-J, Lee E-J, et al. Ubiquitin specific protease 19 involved in transcriptional repression of retinoic acid receptor by stabilizing CORO2A. Oncotarget 2016;7:34759–72.

[58] Mei Y, Hahn AA, Hu S, Yang X. The USP19 deubiquitinase regulates the stability of c-IAP1 and c-IAP2. The Journal of biological chemistry 2011;286:35380–7.

[59] Nakamura N, Harada K, Kato M, Hirose S. Ubiquitin-specific protease 19 regulates the stability of the E3 ubiquitin ligase MARCH6. Experimental cell research 2014;328:207–16.

[60] Wu X, Lei C, Xia T, Zhong X, Yang Q, Shu HB. Regulation of TRIF-mediated innate immune response by K27-linked polyubiquitination and deubiquitination. Nature communications 2019;10:4115.

[61] Altun M, Zhao B, Velasco K, Liu H, Hassink G, Paschke J, et al. Ubiquitin-specific protease 19 (USP19) regulates hypoxia-inducible factor 1alpha (HIF-1alpha) during hypoxia. The Journal of biological chemistry 2012;287:1962–9.

[62] Harada K, Kato M, Nakamura N. USP19-Mediated Deubiquitination Facilitates the Stabilization of HRD1 Ubiquitin Ligase. International journal of molecular sciences 2016;17.

[63] Wu M, Tu H-q, Chang Y, Tan B, Wang G, Zhou J, et al. USP19 deubiquitinates HDAC1/2 to regulate DNA damage repair and control chromosomal stability. Oncotarget 2017;8:2197–208.

[64] Lu Y, Bedard N, Chevalier S, Wing SS. Identification of distinctive patterns of USP19-mediated growth regulation in normal and malignant cells. PLoS One 2011;6:e15936.

[65] Tse WK, Eisenhaber B, Ho SH, Ng Q, Eisenhaber F, Jiang YJ. Genome-wide loss-of-function analysis of deubiquitylating enzymes for zebrafish development. BMC Genomics 2009;10:637.

[66] Farshi P, Deshmukh RR, Nwankwo JO, Arkwright RT, Cvek B, Liu J, et al. Deubiquitinases (DUBs) and DUB inhibitors: a patent review. Expert Opin Ther Pat 2015;25:1191–208.

[67] Fraile JM, Quesada V, Rodriguez D, Freije JM, Lopez-Otin C. Deubiquitinases in cancer: new functions and therapeutic options. Oncogene 2012;31:2373–88.

[68] Luise C, Capra M, Donzelli M, Mazzarol G, Jodice MG, Nuciforo P, et al. An atlas of altered expression of deubiquitinating enzymes in human cancer. PLoS ONE 2011;6.

[69] Wei R, Liu X, Yu W, Yang T, Cai W, Liu J, et al. Deubiquitinases in cancer. Oncotarget 2015;6:12872–89.

[70] D’Arcy P, Wang X, Linder S. Deubiquitinase inhibition as a cancer therapeutic strategy. Pharmacol Ther 2015;147:32–54.

[71] Poondla N, Chandrasekaran AP, Kim K-S, Ramakrishna S. Deubiquitinating enzymes as cancer biomarkers: new therapeutic opportunities? BMB Reports 2019;52:181–9.

[72] Jacq X, Gavory G, O’Dowd C, Cranston A, Baker O, Bell C, et al. Discovery and development of first-in-class orally bioavailable USP19 inhibitors [abstract]. . American Association for Cancer Research Annual Meeting 2019. Atlanta, GA. Philadelphia (PA): Cancer Res 2019 2019.

[73] Li Z, Yin S, Zhang L, Liu W, Chen B. Prognostic value of reduced E-cadherin expression in breast cancer: a meta-analysis. Oncotarget 2017;8:16445–55.

[74] Yang L, Wang XW, Zhu LP, Wang HL, Wang B, Zhao Q, et al. Significance and prognosis of epithelial-cadherin expression in invasive breast carcinoma. Oncol Lett 2018;16:1659–65.

[75] Liu CC, Prior J, Piwnica-Worms D, Bu G. LRP6 overexpression defines a class of breast cancer subtype and is a target for therapy. Proc Natl Acad Sci U S A 2010;107:5136–41.

[76] King TD, Suto MJ, Li Y. The Wnt/beta-catenin signaling pathway: a potential therapeutic target in the treatment of triple negative breast cancer. Journal of cellular biochemistry 2012;113:13–8.

[77] Hu W, Su Y, Fei X, Wang X, Zhang G, Su C, et al. Ubiquitin specific peptidase 19 is a prognostic biomarker and affect the proliferation and migration of clear cell renal cell carcinoma. Oncol Rep 2020;43:1964–74.

[78] Liu Q, Zhao S, Su P, Yu S. Gene and isoform expression signatures associated with tumor stage in kidney renal clear cell carcinoma. BMC Systems Biology 2013;7:1–11.

[79] Sullivan KD, Lewis HC, Hill AA, Pandey A, Jackson LP, Cabral JM, et al. Trisomy 21 consistently activates the interferon response. eLife 2016;5.

[80] Bahnson A, Athanassiou C, Koebler D, Qian L, Shun T, Shields D, et al. Automated measurement of cell motility and proliferation. BMC Cell Biol 2005;6:19.

[81] Tinevez JY, Perry N, Schindelin J, Hoopes GM, Reynolds GD, Laplantine E, et al. TrackMate: An open and extensible platform for single-particle tracking. Methods 2017;115:80–90.

[82] Wiggins H, Rappoport J. An agarose spot assay for chemotactic invasion. Biotechniques 2010;48:121–4.

[83] Salvany L, Muller J, Guccione E, Rorth P. The core and conserved role of MAL is homeostatic regulation of actin levels. Genes & development 2014;28:1048–53.

[84] Ahmed M, Basheer HA, Ayuso JM, Ahmet D, Mazzini M, Patel R, et al. Agarose Spot as a Comparative Method for in situ Analysis of Simultaneous Chemotactic Responses to Multiple Chemokines. Sci Rep 2017;7:1075.

[85] Rameshwar P, Patel S, Pine S, Rameshwar P. Noble Agar Assay for Self-Renewal. Protocol Exchange 2013.

[86] Ampuja M, Jokimäki R, Juuti-Uusitalo K, Rodriguez-Martinez A, Alarmo E-L, Kallioniemi A. BMP4 inhibits the proliferation of breast cancer cells and induces an MMP-dependent migratory phenotype in MDA-MB-231 cells in 3D environment. BMC Cancer 2013;13:1–13.

[87] Farias EF, Petrie K, Leibovitch B, Murtagh J, Chornet MB, Schenk T, et al. Interference with Sin3 function induces epigenetic reprogramming and differentiation in breast cancer cells. Proc Natl Acad Sci U S A 2010;107:11811–6.

[88] Ibrahim AM, Sabet S, El-Ghor AA, Kamel N, Anis SE, Morris JS, et al. Fibulin-2 is required for basement membrane integrity of mammary epithelium. Sci Rep 2018;8:14139.

[89] Sadej R, Romanska H, Baldwin G, Gkirtzimanaki K, Novitskaya V, Filer AD, et al. CD151 regulates tumorigenesis by modulating the communication between tumor cells and endothelium. Molecular cancer research : MCR 2009;7:787–98.

[90] Llorens MC, Rossi FA, Garcia IA, Cooke M, Abba MC, Lopez-Haber C, et al. PKCalpha Modulates Epithelial-to-Mesenchymal Transition and Invasiveness of Breast Cancer Cells Through ZEB1. Front Oncol 2019;9:1323.

[91] Vaske CJ, Benz SC, Sanborn JZ, Earl D, Szeto C, Zhu J, et al. Inference of patient-specific pathway activities from multi-dimensional cancer genomics data using PARADIGM. Bioinformatics 2010;26:i237–45.

[92] Lattanzio R, Marchisio M, La Sorda R, Tinari N, Falasca M, Alberti S, et al. Overexpression of activated phospholipase Cgamma1 is a risk factor for distant metastases in T1-T2, N0 breast cancer patients undergoing adjuvant chemotherapy. Int J Cancer 2013;132:1022–31.

